# Dominance of metabolically flexible fermenters drives intestinal gas production in Crohn’s disease

**DOI:** 10.64898/2026.06.24.733704

**Authors:** Astrid KM Stubbusch, Caitlin Welsh, Lianxin Li, Yuka Katayama, Edward M Giles, Manh Tung Vu, Enes Makalic, Vanessa Rossetto Marcelino, Samuel C Forster, Chris Greening

## Abstract

Molecular hydrogen (H_2_) and hydrogen sulfide (H_2_S) are central gut metabolites that shape microbial metabolism and affect host health. In Crohn’s disease (CD), the shift in microbiota composition (‘dysbiosis’) is associated with intestinal accumulation of these gases, but the responsible microbes remain poorly resolved. Here, we analysed 4,644 bacterial and archaeal species-level genomes from the Unified Human Gastrointestinal Genome Collection to identify H_2_-cycling microbes, assessed their prevalence in ca. 1,700 stool metagenomes from healthy and diseased individuals, and validated their activity using culture-based incubations of stool isolates and biopsy samples. Approximately half of all species encoded H_2_-producing abilities, with acetate- and propionate-forming fermenters such as *Phocaeicola* and *Bacteroides* dominating healthy cohorts, whereas comparatively few taxa, including *Escherichia* and *Megamonas*, encoded H_2_ consuming abilities. In CD, H_2_ producers became more abundant but less diverse, favouring species with multiple H_2_-evolving hydrogenases and more fermentation routes, especially *Clostridium* and *Enterocloster* species. Consistently, isolates enriched in CD produced H_2_ faster and at higher concentrations than health-associated isolates. Increased H_2_S-producing capacity in CD was driven mainly by these H_2_-producing fermenters carrying anaerobic sulfite reductases (Asr), rather than sulfate-reducing bacteria, and was supported by elevated H_2_S production in Asr-positive isolates, likely providing an additional electron sink. These findings provide a species-resolved view of gut gas metabolism and implicate metabolically flexible fermenters in excessive gas and sulfide production in gut disorders.

## Introduction

The gut microbiota contributes to human digestion while also producing substantial amounts of intestinal gas. By fermenting undigested substrates, including dietary fibre and amino acids, intestinal microorganisms produce large quantities of fermentation products - such as short-chain fatty acids (SCFAs; acetate, propionate and butyrate), other organic acids (mainly lactate and succinate), and alcohols - that supply energy to human cells and shape immunity and inflammation^1,2^ **(Fig. 1)**. Fermentation also produces gas as a by–product^3^. In healthy individuals, the gut microbiota produces approximately 0.2-1.5 L of gas per day^4^, consisting mainly of hydrogen (H_2_), carbon dioxide (CO_2_), and methane (CH_4_), and < 1% sulfur–containing trace gases such as hydrogen sulfide (H_2_S)^4,5^. Among these, H_2_ plays a central role in microbial metabolism in the gastrointestinal tract, impacting human nutrition and health^6^. It is formed when excess electrons from fermentation, carried by reduced redox cofactors such as ferredoxin and NADH, are transferred to protons (H⁺) and released into the gut H_2_ pool as diffusible gas **(Fig. 1)**. Roughly one third of the H_2_ is consumed by microbes, whereas the rest is excreted as flatus or via breath^6–8^. Classically, three guilds of H_2_ consumers have been considered: reductive acetogens, sulfate reducing bacteria, and methanogens^4,9^. More recently, ammonifiers, iron reducers, and anaerobic respirers of dietary or host-derived compounds have also been proposed as H_2_ consumers^10^ **(Fig. 1)**. Both H_2_ production and consumption are catalysed by hydrogenases, specialised metalloenzymes that mediate H_2_ production, consumption, and sensing^11^ **(Fig. 1, Table S1)**, with the H_2_-producing group B [FeFe] hydrogenases representing the most widespread hydrogenases in the human gut^12^. However, although intestinal gas levels depend on the balance between microbial H_2_ production and consumption^13^, its microbial mediators are not well understood.

**Fig. 1:**
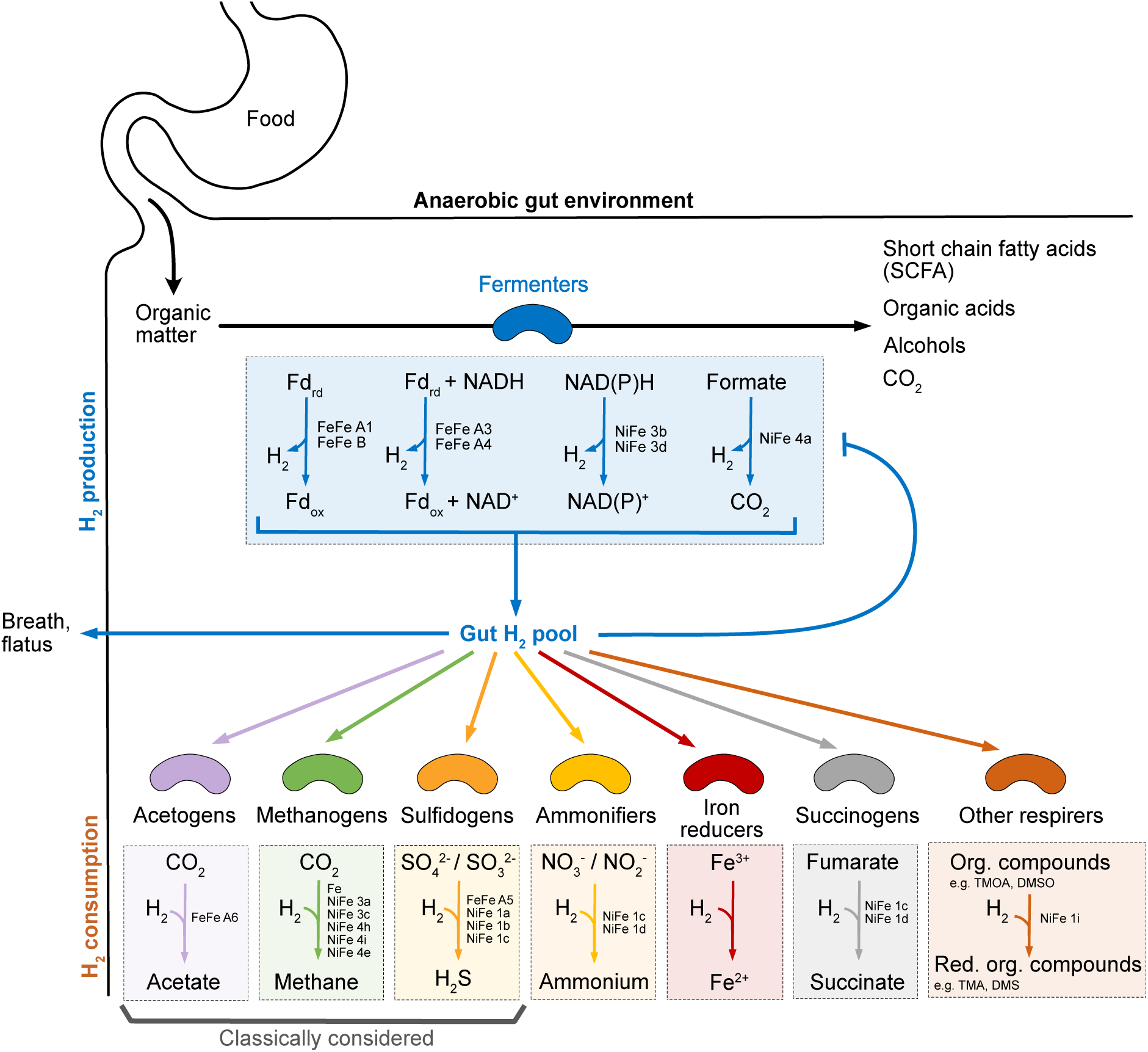
Hydrogen (H_2_) producing and consuming metabolic pathways of microorganisms in the gastrointestinal tract. H_2_ producers and consumers are characterised by specific hydrogenase types and coupled H_2_-producing or -consuming pathways. Org. = Organic; Red. = Reduced. Based on previous reports^34–36^.

Major shifts in the microbial community composition (so-called dysbiosis) are a consistent phenomenon in intestinal inflammatory disorders, such as Inflammatory Bowel Disease (IBD). IBD is frequent in high-income countries and globally increasing^14^. Crohn’s disease (CD) is a major form of IBD, characterized by chronic inflammation of the gastrointestinal tract^15^. Symptoms include bowel urgency, diarrhoea, flatulence, and bloating^16^. Although its precise etiology remains unclear, CD is considered to arise from dysregulated interactions between the gut microbiota and the host immune system^17^. Patients with CD exhibit a consistent gut microbiota dysbiosis, including reduced microbial diversity, depletion of SCFA–producing taxa, and expansion of pro-inflammatory species. This altered community structure is thought to exacerbate intestinal inflammation both by modulating immune responses and by disrupting microbial metabolism, with downstream effects on epithelial cell function and barrier integrity^17^. Despite extensive taxonomic profiling, the functional links between dysbiotic microbial communities and shifts in metabolic output - particularly metabolites such as H_2_, hydrogen sulfide (H_2_S), and SCFAs - remain poorly understood.

Imbalances in microbially-derived gas have been linked to IBD. Excessive H_2_ formation causes discomfort through bloating^18^, thermodynamically inhibits fermentation^13,19^ **(Fig. 1)**, shifting microbial metabolism towards internally redox-balanced fermentation such as ethanol and lactate formation^20,21,6^. In CD, changes in hydrogen metabolism have been suggested by a depletion of stool microbes with the most widespread hydrogenase type^12^, but the consequences for hydrogen metabolism in CD patients remain largely unexplored. CD has also been linked to increased levels of the gas H_2_S, which may contribute to disease etiology^13,22–24^. Microbially produced H_2_S can accumulate to toxic concentrations that impair colonocyte metabolism, damage the intestinal mucosa, and promote gastrointestinal inflammation in a concentration-dependent manner^22,23,25–29^. Gut microbial H_2_S production has classically been attributed to sulfate-reducing bacteria (SRB) such as *Desulfovibrio* and *Bilophila*, which reduce inorganic sulfate or sulfite to H_2_S via dissimilatory sulfate reductases (Dsr) to conserve energy^23,30,13^. However, anaerobic sulfite reductases (Asr) also generate H_2_S from sulfite, derived from both organic substrates (e.g. cysteine) and inorganic substrates (e.g. sulfate/sulfite)^31,32^ (**Fig. S1**). Notably, Asr genes are approximately twice as abundant as Dsr genes in human stool^32^, and Asr-encoding taxa are increasingly considered as contributors to gut H_2_S formation^33,32,13,23,28^, though their metabolic role and drivers remain little understood.

Here, we systematically identified microbes involved in intestinal H_2_ metabolism and H_2_S production at species-level resolution in health and CD, and tested these inferences through culture- and biopsy-based measurements of gas production. We found that about half of gut species encode H_2_-producing pathways, with diverse acetate- and propionate-forming fermenters dominating healthy microbiomes, while H_2_ consumption was restricted to a small number of taxa. In CD, H_2_ producers became less diverse but proportionally more abundant, favouring taxa such as *Clostridium* and *Enterocloster* encoding multiple hydrogenases that exhibited higher H_2_ production rates *in vitro*. Elevated H_2_S-producing capacity in CD was driven primarily by fermentative bacteria encoding anaerobic sulfite reductases (*asr*) rather than classical sulfate-reducing bacteria (SRB). This CD-associated shift towards multi-hydrogenase, *asr*-positive fermenters likely exacerbates both H_2_ accumulation and H_2_S formation, contributing to bloating and inflammation, and directing therapeutic efforts to CD-associated fermenters rather than classically considered sulfate-reducing bacteria.

## Results and discussion

### Half of gut-dwelling prokaryotic species encode hydrogen-evolving capabilities, while few possess hydrogen-consuming capabilities

To identify the bacteria and archaea that control H_2_ levels in the human gut, we developed a framework to identify potential H_2_ producers (hydrogenogens) and H_2_ consumers (hydrogenotrophs) based on their genome and applied it to the Unified Human Gastrointestinal Genome Collection (UHGG)^37^. This genome collection comprises 4,644 gut prokaryotic species, of which ∼30% have cultured representatives and ∼70% were detected by metagenomic sequencing only. First, we identified all hydrogenase enzymes through homology-based searches, then classified them and predicted their function based on sequence phylogeny and gene neighbourhood^36,38,39^ (**Table S1, S2**). In the 4,616 bacterial species, the most common hydrogenases were the ferredoxin-dependent H_2_-evolving group B [FeFe]-hydrogenase (37%) and electron-bifurcating group A3 [FeFe]-hydrogenase (26%).

Also widespread were the functionally uncharacterised group A2 [FeFe]-hydrogenase (17%), the energy-converting group 4e [NiFe]-hydrogenase (15%), and the formate-disproportionating group 4a [NiFe]-hydrogenase (12%) (**Fig. 2A**).

**Fig. 2.**
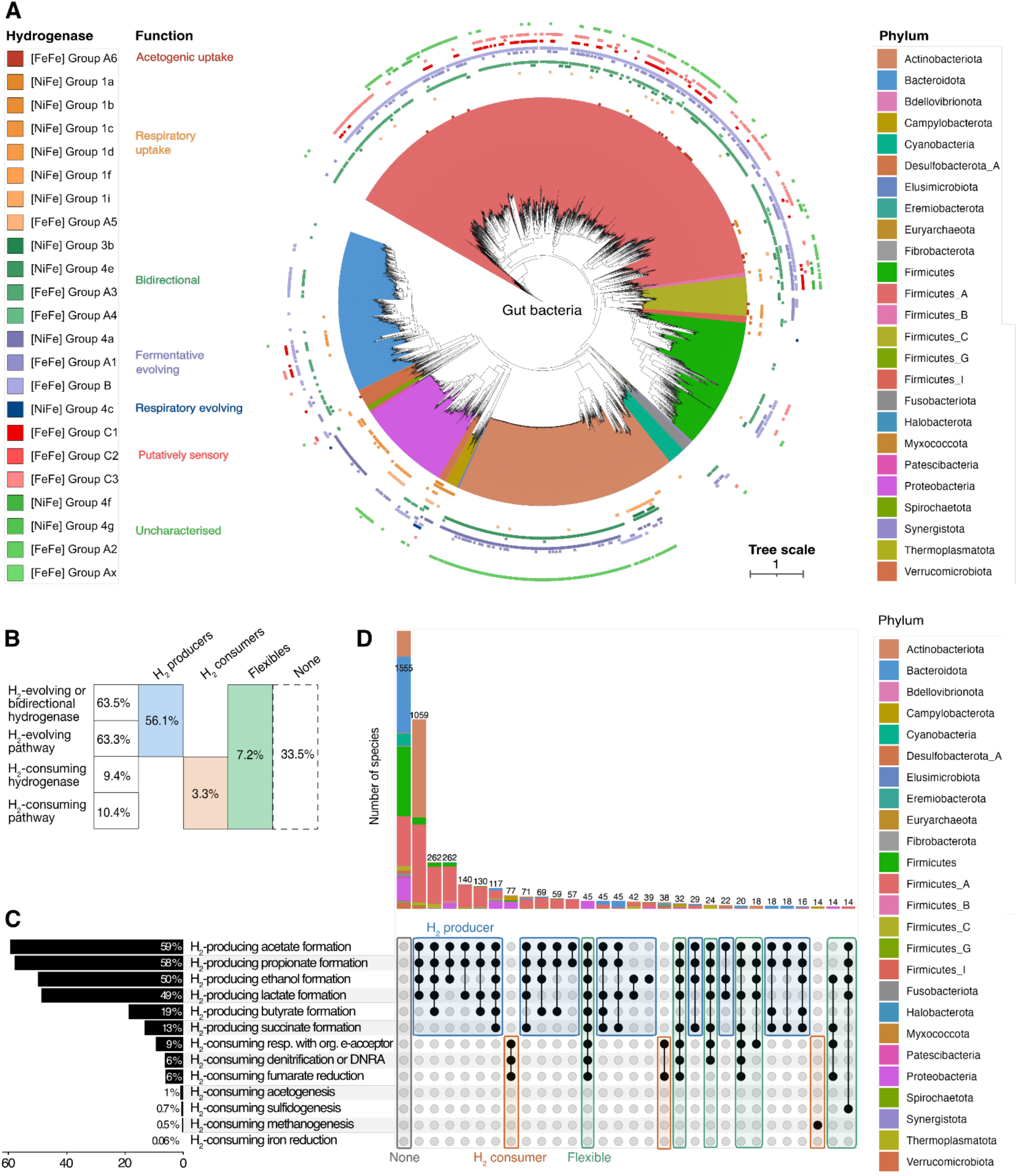
Phylogenetically and metabolically diverse bacteria and archaea of the human gastrointestinal tract can cycle H_2_. **(A)** Phylogenetic distribution of hydrogenase types and associated function in bacteria of the Unified Human Gastrointestinal Genome Collection (UHGG). Hydrogenases were widespread, especially the fermentative H_2_-evolving group B [FeFe] hydrogenase (37% of species) and the bidirectional group A3 [FeFe] hydrogenase (26% of species). Nomenclature based on GTDB. **(B)** Schematic of the prediction of potential H_2_ producers (blue), H_2_ consumers (orange), flexibles (green), and non-H_2_-cyclers (grey), alongside the respective fractions of bacterial and archaeal species with these capabilities. **(C)** H_2_-cycling pathways utilized by H_2_ producers and H_2_ consumers, respectively. The fractions show the number of UHGG species that possess potentially H_2_ producing or consuming pathways in combination with a H_2_ evolving or H_2_ consuming hydrogenase, respectively. **(D)** Number (bars) and taxonomy (colour) of UHGG species with the same set of H_2_-producing and H_2_-consuming capabilities. The upset plot marks the presence (black dot) and absence (no dot) of a potentially H_2_-producing or consuming metabolic pathway alongside a H_2_-evolving or consuming hydrogenase, respectively. Capability sets that characterise potential H_2_ producers are marked in blue, H_2_ consumers in orange, flexibles with both characteristics in green, and non-H_2_-cyclers with neither characteristic in grey. This colour scheme and definition is consistent with (B).

Methanogenesis-associated [NiFe]- and [Fe]-hydrogenases were also encoded in 25 of the 28 archaea species of the human gut (89%), sometimes accompanied by the bidirectional group 4e [NiFe] hydrogenase (32%) (**Fig. S2**).

To identify likely H_2_ producers and consumers, we further assessed whether the presence of H_2_-evolving hydrogenases was paired with the presence of potential H_2_-producing fermentation pathways, and the presence of H_2_-consuming hydrogenases with potential H_2_-consuming metabolic pathways. In the 4,644 bacterial and archaeal species found in the human gut, we identified 2,604 potential H_2_ producers (56.1%), 151 potential consumers (3.3%), 334 potential flexibles (7.2%) that have both H_2_-producing and H_2_-consuming capabilities, and 1,555 non-H_2_-cyclers (33.5%) in which no hydrogenase with associated pathway for H_2_ production or consumption was found (**Fig. 2B**). Non-H_2_-cyclers that lack hydrogenases may still carry out fermentation, which they redox-balanced internally or through other means than H_2_ production. Overall, Bacillota (former Firmicutes), Spirochaetota, and Actinomycetota (former Actinobacteriota) had the highest fraction of potential H_2_ producers (**Table S3**). The phyla with the highest fraction of potential H_2_ consumers were the archaeal Methanobacteriota (former Euryarchaeota) and Thermoplasmatota, and Campylobacterota (82%, 36%, 51% of species, respectively, **Table S3**).

To understand which H_2_ producing and consuming pathways are most common in prokaryotic species of the human gut, we assessed their presence and their common combinations within gut microbes. We found that most prevalent H_2_-producing pathways in H_2_ producers were fermentative acetate and propionate formation, encoded in nearly 60% of the prokaryotic species found in the human gut **(Fig. 2C)**. The combinations of possible H_2_ producing pathways were quite diverse in potential H_2_ producers, with phylogenetically conserved preferences for certain combinations of potentially H_2_ producing pathways (**Fig. 2D**). Most commonly, H_2_ producers were capable of acetate and propionate formation, often in combination with further fermentation pathways. The fermentation strategies of H_2_ producers differed markedly across microbial phyla. While acetate formation emerged as the dominant H_2_-generating pathway in Bacillota (former Firmicutes), Actinomycetota (former Actinobacteriota), and Bacteroidota (former Bacteroidetes), butyrate formation was the prominent H_2_-forming pathway in the phyla Fusobacteriota, Synergistota, and Bacillota (**Table S4**).

For H_2_-consuming pathways, anaerobic respiration using dietary and host-derived organic electron acceptors including fumarate was most prevalent, encoded by approximately 9% of all species and far more prevalent than classical hydrogenotrophic pathways, including acetogenesis, sulfidogenesis, and methanogenesis (Fig. 2C). Overall, H_2_ consumers most commonly possessed genes for both organic electron acceptors for anaerobic respiration and nitrate reduction (DNRA and/or denitrification steps), e.g. in the phyla Pseudomonadota (former Proteobacteria), Campylobacterota, and Desulfobacterota, followed by methanogens (**Fig. 2D, Table S4**).

### H_2_ producers are more abundant than H_2_ consumers in the human gut

To determine the abundance of H_2_ producers and consumers in the human gut, we assessed their abundance within stool samples from 888 healthy individuals^24^. We found a median abundance of 76% H_2_ producers, 0.5% H_2_ consumers, 1.2% flexibles, and 19% non-H_2_-cyclers across all individuals (**Fig. 3A**). Consistent with genome-catalog predictions **(Fig. 2C and D)**, the most abundant H_2_-cycling pathways in stool samples, defined by the abundance of cells encoding them, were fermentative co-associated acetate and propionate production in H_2_ producers, and reduction of organic compounds such as fumarate in H_2_ consumers **(Fig. 3B)**.

**Fig. 3.**
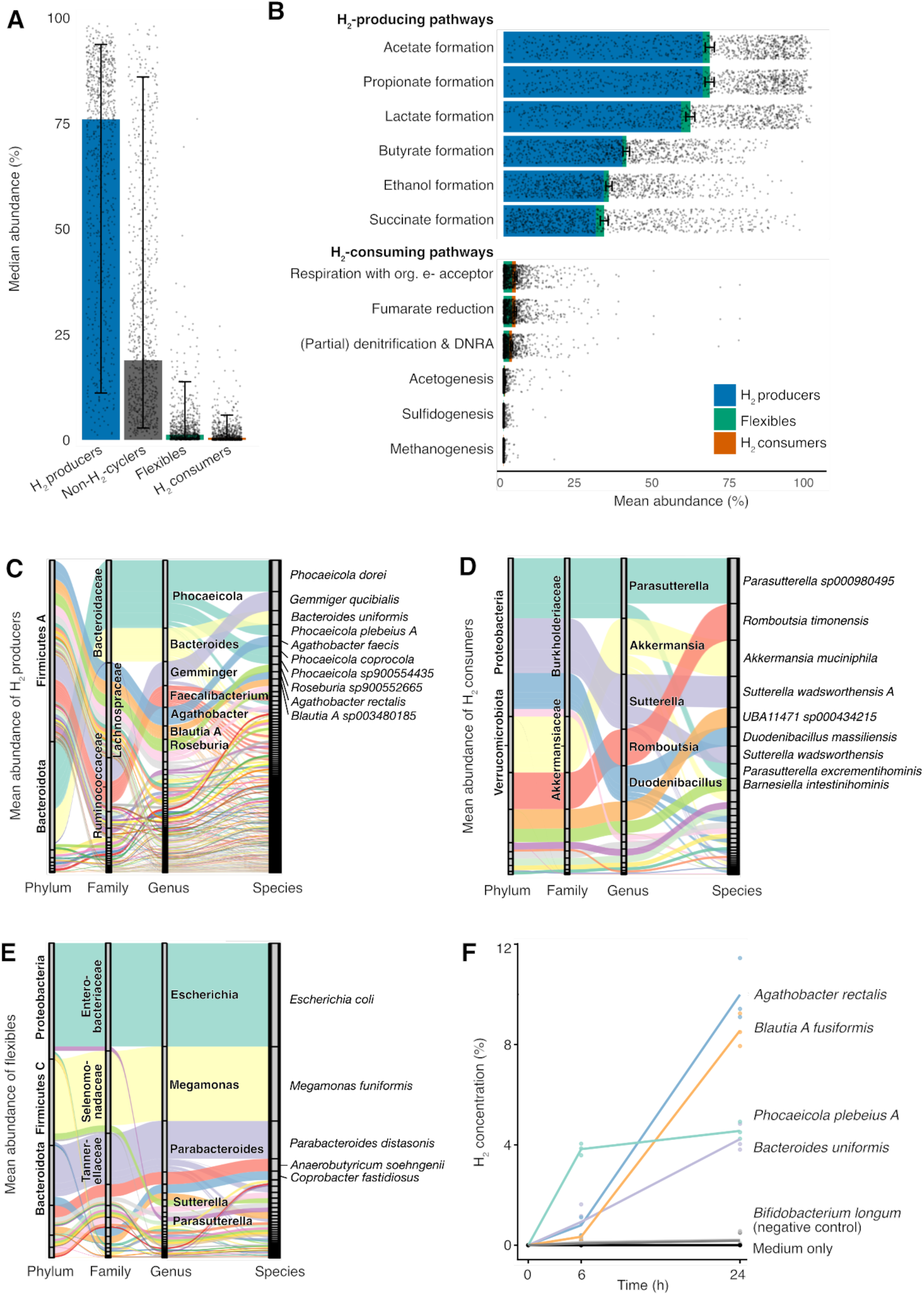
H_2_ producers are more abundant and phylogenetically diverse than H_2_ consumers in stool samples from 888 healthy individuals. **(A)** Median abundance of predicted H_2_ producers (blue), non-H_2_-cyclers (grey), flexibles (green), and H_2_ consumers (orange) in stool samples of 888 healthy individuals. Error bars show the 90% confidence interval for the median. **(B)** Mean abundance of H_2_-producing and H_2_-consuming capabilities, i.e. H_2_-producing and H_2_-consuming metabolic pathways paired with a respective hydrogenase type. Dots indicate the abundance in each sample, and error bars show the 90% confidence interval of the mean. **(C)** Mean abundance and taxonomic identity of the H_2_ producers. **(D)** Mean abundance and taxonomic identity of the H_2_ consumers. **(E)** Mean abundance and taxonomic identity of the flexibles that harbour respective hydrogenases and pathways to either produce or consume H_2_. **(F)** Gas chromatographic measurement of H_2_ production by predicted H_2_ producers over time growing in YCFA broth. Dots show three biological replicates; lines connect the mean of the triplicates.

We could resolve the identity of the H_2_ producers and consumers from these stool samples on a species-resolved level (**Table S5**). The most abundant H_2_ producers in stool were *Phocaeicola dorei* (7% ± 8%, mean ± s.d. across all metagenomes), *Gemmiger qucibialis* (4% ± 7%), and *Bacteroides uniformis* (3% ± 4%) (**Fig. 3C**, **Tables S6**). The fermentation routes of H_2_ producers varied by taxon: while *Phocaeicola* and *Bacteroides* were the most abundant predicted producers of acetate, propionate, lactate, and succinate, *Phocaeicola*, *Gemmiger, Faecalibacterium,* and *Agathobacter* predominated in predicted butyrate formation, and *Phocaeicola, Blautia A,* and *Lachnospira* in ethanol formation (**Fig. S4**).

The dominant H_2_ consumers were *Parasutterella sp000980495* (0.2% ± 1.0%, mean ± s.d. of relative abundance), *Romboutsia timonensis* (0.2% ± 1.3%), *Akkermansia muciniphila* (0.2% ± 0.7%), and *Sutterella wadsworthensis A* (0.1% ± 0.8%) (**Fig. 3D, Table S7**). Dominating flexibles, capable of both H_2_ production and consumption, were *Escherichia coli* (1% ± 4.6%), *Megamonas funiformis* (0.7% ± 2.9%), and *Parabacteroides distasonis* (0.4% ± 0.9%) (**Fig. 3E, Table S8**). Prevalent H_2_ consumption pathways varied by taxon: *Escherichia, Megamonas*, and *Parabacteroides* predominated in respiration with organic electron acceptors (e.g fumarate reduction); *Escherichia, Parabacteroides*, and *Sutterella* in nitrate/nitrite reduction, *Romboutsia* and *Anaerobutyricum* in acetogenesis; *Bilophila* and *Desulfovibrio* in sulfidogenesis; and the archaeal genera *UBA71*, *Methanobrevibacter*, and *Methanomethylophilus* in methanogenesis (**Fig. S5**).

To test whether the taxa that we predict to be the most abundant mediators of H_2_ production are indeed acting as H_2_ producers, we cultured stool-derived isolates in YCFA broth and measured accumulating H_2_ in the headspace of the culturing vials after 0, 6, and 24 hours by gas chromatography. We found that *Bacteroides uniformis* (relative abundance 2.9% ± 0.8%, mean ± s.d.), *Phocaeicola plebeius A* (2.4% ± 6.8%), and *Agathobacter rectalis* (1.5% ± 2.4%) produced H_2_ during growth, reaching H_2_ concentrations of approximately 4% to 10%, while no H_2_ production was measured for the hydrogenase-lacking^12^ negative control *Bifidobacterium longum* (**Fig. 3F, Table S9)**. *Blautia A fusiformis* (relative abundance 1.5% ± 2.9% of *Blautia A sp003480185*) also produced H_2_, as predicted from its ferredoxin-dependent H_2_-evolving group A1 and B [FeFe]-hydrogenases. Although some *Blautia* species such as *Blautia hydrogenotrophica* are well-known hydrogenotrophic acetogens^6,40^, the absence of genes encoding acetogenic uptake hydrogenases in this *Blautia* genome suggests that some *Blautia* are unlikely to function as H_2_-dependent acetogens. This may be related to previously observed variation in the Wood-Ljungdahl pathway^41,42^, in which *Blautia* species do not use H_2_ and CO_2_ for acetogenesis^43^, highlighting a versatile role of acetogens in gut H_2_-cycling. The predicted H_2_ production by *Phocaeicola dorei* (6.5% ± 8.5%) and *Phocaeicola plebeius* (1.0% ± 2.6%) has previously been experimentally shown^12^. These findings confirmed H_2_ production during growth by several of the predicted H_2_ producers that were most abundant in the stool of healthy individuals.

### Increased fraction of H_2_ producers with reduced diversity in CD dysbiosis

To understand whether H_2_ cyclers were affected by gastrointestinal diseases, we compared their abundance in stool samples from healthy individuals and Crohn’s Disease (CD) patients, which had been assembled from several studies^24^. We found a significant increase in the proportion of H_2_ producers (7% increase from a median abundance of 76% in healthy patients) and flexibles (3% increase from a median abundance of 1% in healthy patients) (**Fig. 4A**). As a result, the balance of H_2_ producers to H_2_ consumers and flexibles was significantly decreased (mean log2-ratio of 4.6 in healthy and 3.4 in CD, *p* = 2×10^−5^). This ‘dysbalance’ of H_2_ cyclers was observed across several disorders, but was particularly pronounced in CD **(Fig. S6)**.

**Fig. 4.**
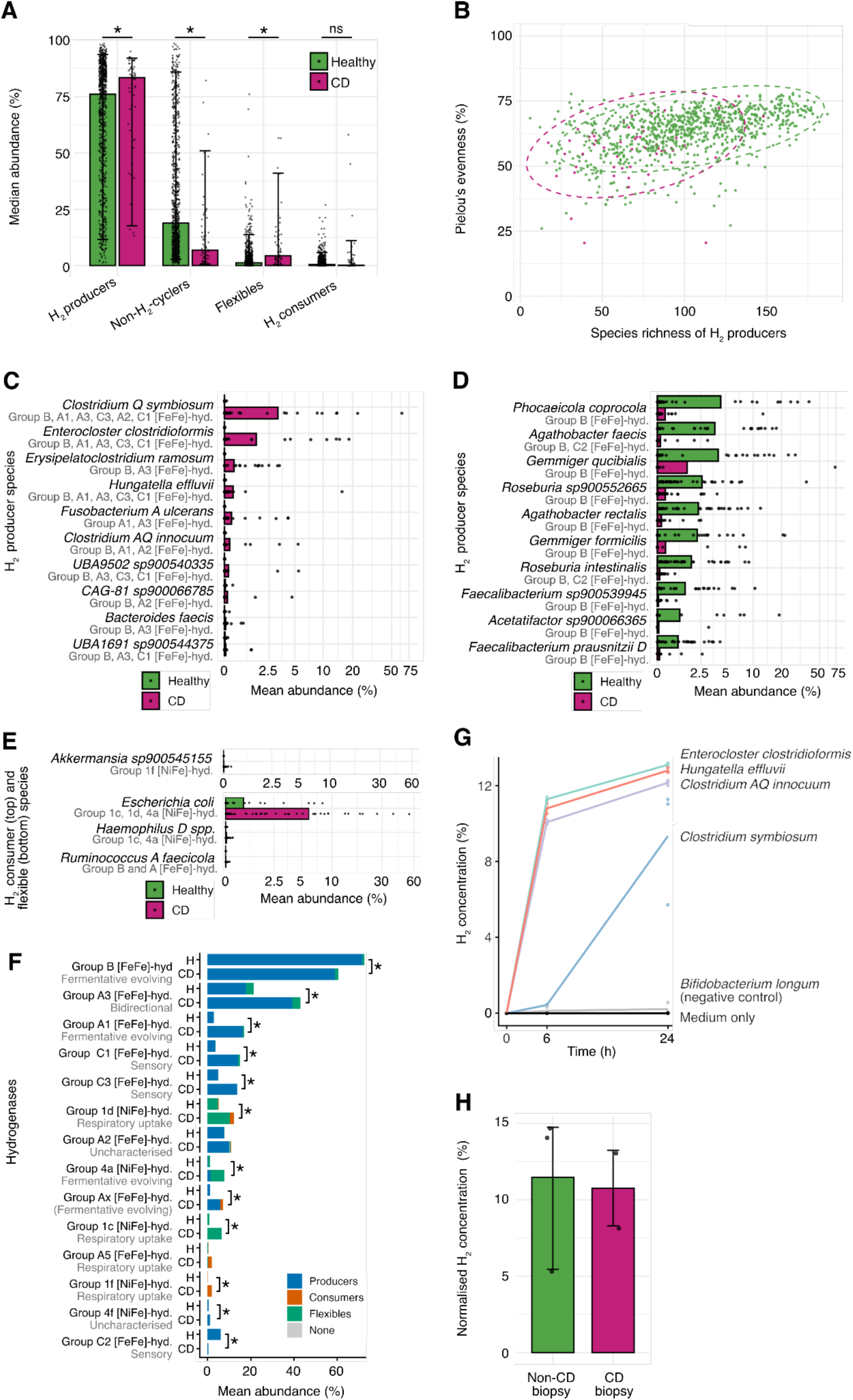
Shifts in mediators of H_2_ cycling and their hydrogenases in CD patients. **(A)** Relative abundance of H_2_ cyclers in healthy and CD patients. Median abundances with 90% CI are shown (n healthy = 888, n CD = 59; Wilcoxon rank-sum test, *p*-values for H_2_ producers 0.03, non-H_2_-cyclers 2×10^−8^, flexibles 10^−11^, H_2_ consumers 0.1). **(B)** Decrease of species richness (x-axis) and Pielou’s evenness index (y-axis) in Crohn’s disease patients compared to healthy individuals. **(C)** The 10 H_2_ producer species with the most increased abundance in CD patients. Ordered by maximal difference in mean abundance in CD patients and healthy controls (n healthy = 39, n CD = 46; Wilcoxon rank-sum test, BH-adjusted *p* < 0.05; MaAsLin *p* < 0.05). **(D)** The 10 H_2_ producer species with the most decreased abundance in CD patients. Ordered by maximal difference in mean abundance in CD patients and healthy controls (n healthy = 39, n CD = 46; Wilcoxon rank-sum test, BH-adjusted *p* < 0.05; MaAsLin *p* < 0.05). **(E)** The one H_2_ consumers species with significantly increased abundance in CD patients. Ordered by maximal difference in mean abundance in CD patients and healthy controls (n healthy = 39, n CD = 46; Wilcoxon rank-sum test, BH-adjusted *p* < 0.05; MaAsLin *p* < 0.05). **(F)** Abundance of taxa with certain hydrogenase types in CD patients and healthy controls. Hydrogenase types of less than 1% abundance in either healthy or CD patients were not shown (n healthy = 39, n CD = 46; Wilcoxon rank-sum test, *p*-values 0.02 (FeFe Group B), 3×10^−5^ (FeFe Group A3), 10^−7^ (FeFe Group A1), 2×10^−5^ (FeFe Group C1), 0.002 (FeFe Group C3), 0.0004 (NiFe Group 1d), 0.8 (FeFe Group A2), 8×10^−6^ (NiFe Group 4a), 0.003 (FeFe Group Ax), 2×10^−6^ (NiFe Group 1c), 0.09 (FeFe Group A5), 0.005 (NiFe Group 1f), 0.04 (NiFe Group 4f), 3×10^−12^ (FeFe Group C2)). H = healthy, CD = Crohn’s disease. **(G)** Gas chromatographic measurement of H_2_ production by predicted H_2_ producers with increased abundance in CD patients over time growing in YCFA broth. Dots show three biological replicates, lines connect the means of the triplicates. **(H)** H_2_ production of biopsy samples from patients diagnosed as CD patients or non-CD patients after 48 h in YCFA medium, normalized by subtracting the media control.

Despite the increased relative abundance of H_2_ producers in CD, their species richness and evenness decreased (**Fig. 4B**), indicating dominance of fewer H_2_ producing species in CD patients (richness ± s.d.: 102 ± 37 in healthy to 72 ± 31 in CD patients, Pielou’s evenness index ± s.d.: 0.63 ± 0.08 in healthy to 0.57 ± 0.11 in CD patients). The species richness of H_2_ flexibles increased, while the number of H_2_ consumer species remained similar (**Fig. S7)** (5.9 ± 2.5 to 7.5 ± 2.8 in flexibles, 2.8 ± 1.9 to 2.1 ± 2.0 in H_2_ consumers). Observed differences were maintained when focusing on a single study of CD patients and healthy case-controls (**Supplementary Note 1**). Overall, the data suggest a shift toward dominance by a few highly abundant H_2_-producing species in CD, while H_2_-flexible taxa became more diverse.

### Dominant H_2_ producers in CD patients encode multiple hydrogenases and show elevated H_2_ production

To reveal the dominant and depleted taxa in CD, we computed the differential abundance of each species between CD patients and healthy case-controls (**Table S10, S11**). Significant abundance changes were identified by both Wilcoxon rank-sum test and fitted multivariable linear models that accounted for compositional effects and for age and BMI of the individuals. We found 11 significantly increased H_2_ producer species in CD patients, especially *Clostridium Q symbiosum* and *Enterocloster clostridioformis* (**Fig. 4C, Table S12**), while 55 H_2_ producer species were significantly depleted (**Fig. 4D, Table S13**) (*p* < 0.05). In addition, one putatively H_2_-consuming *Akkermansia* species was significantly increased in relative abundance in CD patients, as well as three flexible species, including predominantly *Escherichia coli* (**Fig. 4E, Table S14, S15**). Enrichment analysis revealed that H_2_ producers were overrepresented among the species that declined (**Supplementary Note 2**), highlighting that many H_2_-producing species were depleted while a few dominant taxa became more prevalent.

We hypothesized that the dominant H_2_ producers possess different hydrogenases, as a previous gene-centric study had shown a depletion of group B [FeFe]-hydrogenases in favour of other fermentative hydrogenases in stool of CD patients^12^. Overall, H_2_ producers in stool from CD patients possessed on average 2.5 hydrogenases, which was significantly more than healthy controls with 1.7 hydrogenases (Wilcoxon rank-sum test, *p* = 0.03) (**Fig. S9**). H_2_ producers with the fermentative H_2_-evolving group B [FeFe]-hydrogenase, which drives most fermentative H_2_ production in the healthy human gut^12^, and the sensory group C2 [FeFe]-hydrogenase were significantly decreased in relative abundance in CD (**Fig. 4F, S10A**). Instead, H_2_ producers with group A3 and A1 [FeFe]-hydrogenases and putatively sensory group C1 and C3 [FeFe]-hydrogenases were significantly increased. This trend also held true for the species that were most increased in CD patients (**Fig. 4C**) and most decreased in CD patients (**Fig. 4D**). Encoding multiple hydrogenases can enable organisms to maintain fermentation under different hydrogen concentrations, albeit potentially at reduced ATP synthesis^34,36,44^. For example the saccharolytic rumen bacterium *Ruminococcus albus* switches from using electron-bifurcating group A3 hydrogenases at low H_2_ concentrations (producing 4 ATP) to ferredoxin-dependent group A1 hydrogenase at high H_2_ concentrations (producing only 3.3 ATP) by sensing and responding to H_2_ partial pressures with group C hydrogenases^45,36^. Thus, the relative increase of H_2_ producers with diverse hydrogenases in stool from CD patients suggests a change in the microbiota towards H_2_ producers that can also ferment under elevated H_2_ concentrations.

To test whether these alternative hydrogenases increased the levels of microbially produced H_2_, we cultured isolates of the most CD-enriched fermenters in YCFA broth and measured accumulating H_2_ in the headspace of the culturing vials after 0, 6, and 24 hours by gas chromatography (**Fig. 4G, Table S16**). These CD-enriched taxa with diverse hydrogenases produced significantly higher concentrations of H_2_ both after 6 h and 24 h (**Fig. 4G**) compared to the tested H_2_ producers that were most abundant in healthy individuals (**Fig. 3F**) (Wilcoxon rank-sum test, 6 h: *p* = 2×10^−3^, 24 h: *p* = 5×10^−5^), of which most possessed only group B [FeFe]-hydrogenases (**Table S17**). To test for increased H_2_ production in the gut of CD patients, we retrieved biopsy samples from both CD and non-CD patients, transferred them into YCFA broth for anaerobic culturing, and measured their H_2_ production over time. H_2_ accumulated to high levels in biopsy cultures from CD and non-CD donors. However, no significant difference was observed between the two groups (**Fig. 4H, Table S18**). This may be due to high variation in pre-procedural bowel preparation and inter-personal variation, or potentially increased H_2_ consumption may offset increased H_2_ production in CD biopsy cultures where hydrogenotrophs such as *E. coli* are particularly enriched^46^. More extensive sampling, taking into account high interindividual variability, is needed to test this. Overall, these results suggest that the dominating H_2_ producers in CD patients were fermenting taxa with additional group A and group C hydrogenases, which produced H_2_ more rapidly and to higher concentrations.

### Increased metabolic flexibility enables growth at high H_2_ concentrations in CD

Investigating the encoded fermentation pathways in H_2_ producers revealed that the number of fermentation routes, which lead to different fermentation products, was significantly increased in CD patient stool samples compared to healthy case-controls (**Fig. S11A**). Consistently, grouping fermenting bacteria by their encoded fermentation routes showed that taxa capable of producing a wider diversity of fermentation products were significantly enriched, while those with a more limited product spectrum were decreased **(Fig. 5A)**. This suggested an increase in metabolically flexible fermenters with diverse fermentation routes in CD patients. The fermentation routes that were significantly more abundant within the H2 producers in CD patients were ethanol and lactate formation (**Fig. S11C**). As excessive H_2_ accumulation is known to shift microbial metabolism towards internally redox-balanced fermentation such as ethanol and lactate formation^20,21,6^, this finding suggests a community-wide metabolic shift in adaptation to increased H_2_ accumulation in the colon of CD patients, which may reduce production of SCFAs in favour of ethanol and lactate.

**Fig. 5.**
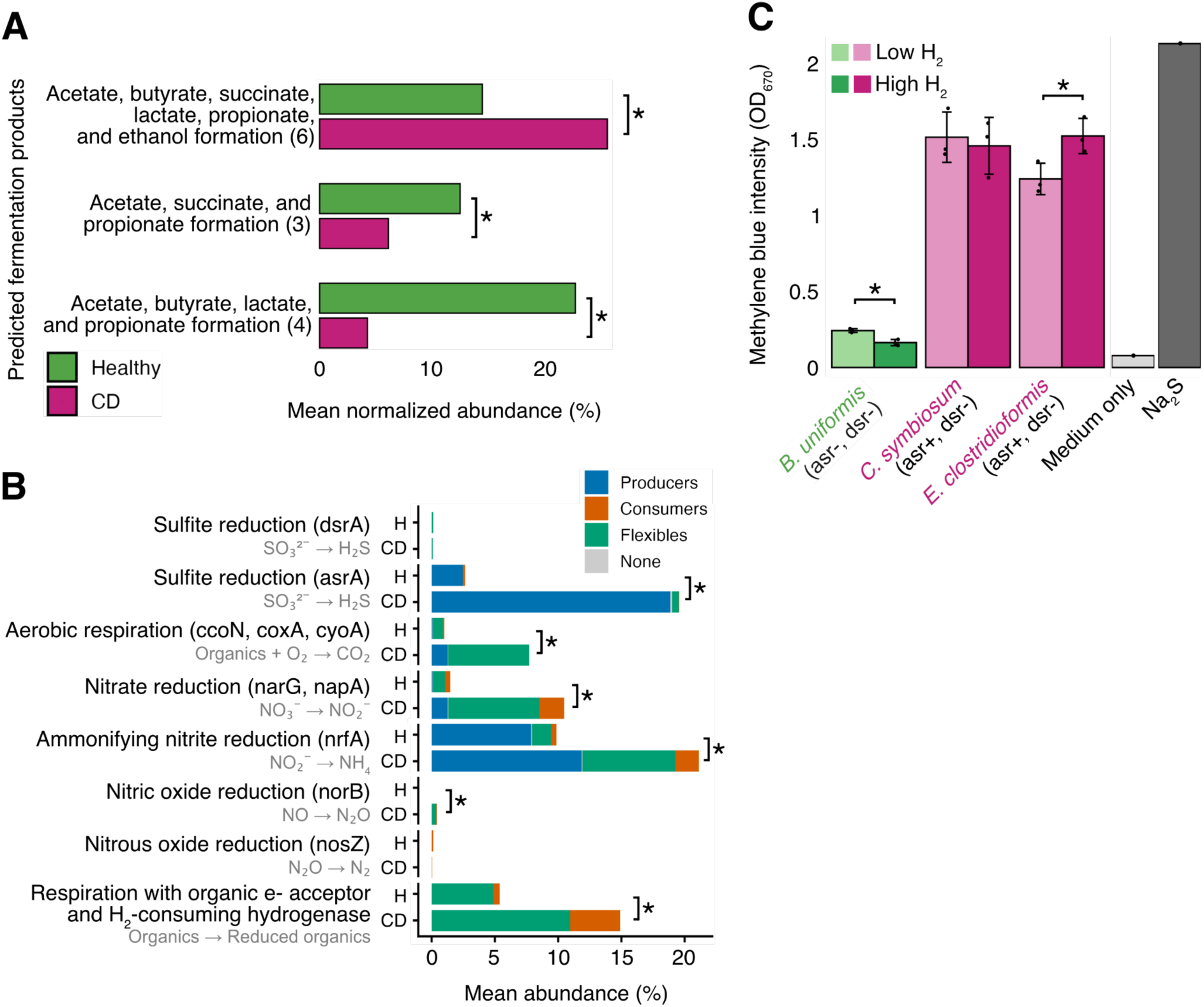
Metabolic flexibility of H_2_ producers in Crohn’s disease. **(A)** H_2_ producers with more predicted fermentation routes were enriched in CD patients, whereas those with fewer possible fermentation products were depleted. Taxa were grouped by shared predicted fermentation products, and only groups with >5% abundance in either condition and an absolute difference >5% between conditions are shown (n_Healthy_ = 39, n_CD_ = 47; Wilcoxon rank-sum test, p-values = 2×10^−2^, 2×10^−3^, 10^−12^, from top to bottom). **(B)** Abundance of taxa with genes linked to H_2_S production, aerobic respiration, denitrification, and DNRA, which have been linked to inflammation in CD (n_Healthy_= 39, n_CD_ = 46; Wilcoxon rank-sum test, *p*-values 0.9 (sulfite reduction with Dsr), 3×10^−7^ (sulfite reduction with Asr), 2×10^−6^ (aerobic respiration), 4×10^−8^ (nitrate reduction), 0.001 (nitrite reduction to ammonia), 2×10^−6^ (nitric oxide reduction), 0.9 (nitrous oxide reduction), 6×10^−5^ (respiration with organic electron acceptors in genomes with a H_2_-consuming hydrogenase). H = healthy, CD = Crohn’s disease. **(C)** H_2_S formation during fermentation under high H_2_ (100% H_2_ atmosphere) and low H_2_ (100% N_2_ atmosphere) by the CD-enriched, *asr*-positive H_2_ producers *C. symbiosum* and *E. clostridioformis*, and the CD-depleted, *asr*-negative *Bacteroides uniformis* (n = 3; *t*-test, p-values = 8×10^−3^, 0.7, 0.03, from left to right). Cell-free medium served as negative control, 200 µM Na_2_S served as positive control.

Also organisms capable of H_2_-dependent respiration were significantly increased in CD patients. Taxa with nitrate reductases (Nar and Nap) and nitrite reductases (Nrf) increased significantly in CD patients (1.5% to 10.5%, Wilcoxon rank-sum test, BH-adjusted *p =* 4×10^−8^; 9.9% to 21.1%, BH-adjusted *p* = 0.001, respectively) (**Fig. 5B**), predominantly *E. coli* with its group 1d and 1c [NiFe]-hydrogenase for respiratory H_2_ uptake (**Fig. S10B**). The capacity for respiration with organic electron acceptors was also significantly increased, mainly through *E. coli* and *Akkermansia muciniphila B* with its group 1f [NiFe]-hydrogenase. The fraction of dissimilatory sulfate reducers remained small. *E. coli* was also capable of aerobic respiration, contributing to a significant increase in capacity for organic respiration (1.0% to 7.7%, BH-adjusted *p* = 2×10^−6^) (**Fig. 5B, S10B**). H_2_-consuming respiration has previously been shown to increase fitness of *E. coli* during gut inflammation, highlighting that respiration of inflammation-derived electron acceptors can boost the growth of commensals like *E. _coli_*_47,48._

While sulfate-reducing bacteria with dissimilatory sulfite reductase gene *dsrA* remained low in abundance (mean 0.1% in both health and CD, BH-adjusted *p* = 0.9), species with an anaerobic sulfite reductase gene *asrA* were significantly increased from a mean abundance of 3% in healthy individuals to 20% in CD patients (BH-adjusted *p* = 3×10^−7^) **(Fig. 5B**). The AsrA-encoding taxa were mostly H_2_ producers, predominantly *Fusobacterium A mortiferum, Clostridium Q symbiosum, Flavonifractor plautii, and Enterocloster clostridioformis* **(Fig. S10B)**, which lacked *dsrA*. Asr regenerates NAD^+^ from NADH^49^ and is regulated by NAD^+^ availability^31,49^, potentially - like H_2_ production - serving as an electron sink for redox-balancing during fermentation.

To test whether Asr supports redox balancing when H_2_ production is thermodynamically unfavorable, we measured H_2_S formation under high H_2_ conditions and, for comparison, low H_2_ conditions. We used YCFA cultures of the *asrA*-positive *C. symbiosum* and *E. clostridioformis*, both H_2_ producers enriched in CD patients **(Fig. 4C and H)**, and the *asrA*-negative *B. uniformis*, a H_2_ producer enriched in healthy controls **(Fig. 3C and F)**. Using the methylene blue assay as a standard colorimetric method for measuring H_2_S after 12 hours, we found that the CD-enriched *C. symbiosum* and *E. clostridioformis* produced high levels of H_2_S, which were further elevated under high H_2_ conditions for *E. clostridioformis* **(Fig. 5C)**. This suggests that H_2_S production may also be used under low H_2_ conditions as an electron sink by *asrA*-positive isolates, and can be further increased in certain isolates by high H_2_ conditions. In contrast, *B. uniformis* produced lower levels of H_2_S, potentially linked to cysteine biosynthesis or degradation^13^ (**Fig. S1**), independent of Asr **(Fig. 5C)**. Together, these results suggest that Asr- and multi-hydrogenase encoding fermenters produce more H_2_S than Asr-lacking fermenters, likely using sulfite reduction as an additional electron sink to sustain fermentation, even under high H_2_ conditions. Given that ∼20% of bacterial cells in stool from CD patients encode Asr, this mechanism may drive H_2_S to toxic concentrations associated with CD pathogenesis^25,28,29,23,50,51^.

## Conclusion

This study provides the first species-resolved view of H_2_ metabolism in the human gut and reveals major shifts in H_2_-producing and H_2_-consuming populations in Crohn’s disease (CD). Across the gut microbiota, H_2_-producing fermenters greatly outnumbered H_2_-consuming respirers, consistent with measurable H_2_ loss by breath and flatus and with a nutrient-rich but electron-acceptor-limited gut lumen that favours fermentation over anaerobic respiration^52^. Notably, our metagenomic analysis indicates that H_2_ consumption linked to respiration with dietary or host-derived electron acceptors, like fumarate, and dissimilatory nitrate reduction to ammonium (DNRA) is more prevalent than the classically emphasized hydrogenotrophic acetogenesis, sulfidogenesis, or methanogenesis^6,53^, redefining the major H_2_ sinks in the human gut.

In CD, the microbiota shifted towards metabolically flexible fermenters, consistent with adaptation to elevated H_2_. CD-associated communities were enriched in taxa encoding multiple H_2_-producing and H_2_-sensing hydrogenases, and isolates of these taxa produced H_2_ more rapidly and to higher concentrations in vitro, consistent with the high prevalence of bloating and flatulence in inflammatory bowel disease^16^. This increased H_2_ production may be regulated by alternative H_2_ sensing hydrogenases ([FeFe] group C1 and C3, rather than C2) and dynamic switching from ferredoxin-dependent group B hydrogenases during H_2_ homeostasis to ferredoxin-dependent group A1 and electron-bifurcating group A3 [FeFe]-hydrogenases under H_2_ backpressure. These CD-enriched taxa also encoded expanded fermentation routes, particularly to ethanol and lactate, suggesting adaptation to high-H_2_ conditions^20,21,6^. We further show that CD-enriched fermenters frequently harbour anaerobic sulfite reductases (Asr) and isolates produced elevated H_2_S in vitro. Sulfite reduction may therefore provide an alternative electron sink when H_2_ production becomes thermodynamically constrained and O_2_-sensitive [FeFe]-hydrogenases are inhibited under inflammatory conditions (**Fig. 6**). This identifies fermentation-driven sulfite reduction, rather than classical dissimilatory sulfate reduction by sulfate-reducing bacteria, as a potential major source of H_2_S in CD.

**Fig. 6.**
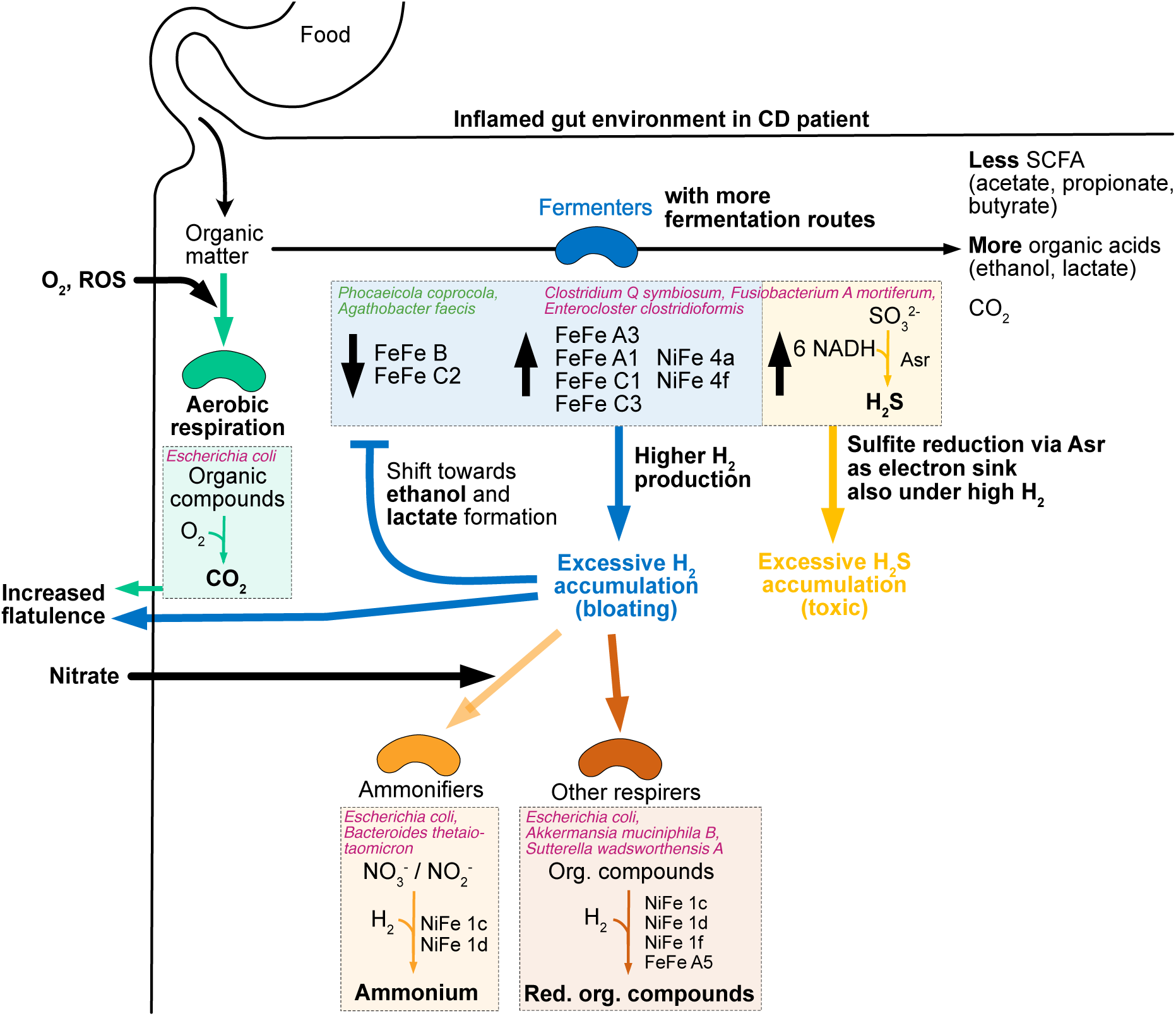
Conceptual model of altered microbial gas metabolism in the inflamed gut environment of CD patients. Metabolically flexible fermenters with (i) more fermentation routes, (ii) alternative fermentative hydrogenases, and (iii) often anaerobic sulfite reductases (Asr) for redox-balancing dominated over fermenters with only group B (and C2) [FeFe]-hydrogenases, which get thermodynamically inhibited by H_2_ accumulation. Fermentation profiles shifted towards ethanol and lactate formation, which require less external redox-balancing through e.g. H_2_ formation than the other fermentation routes. Excessive accumulating H_2_ leads to bloating and increased flatulence. Increased aerobic respiration through inflammation-derived O_2_ and reactive oxygen species (ROS) can lead to additional CO_2_ formation. Accumulating H_2_ and host-derived nitrate, a by-product of the inflammation, can be metabolized to ammonium by the increased fraction of ammonifiers, while other respirers consume H_2_ for respiration with diet- or host-derived organic electron acceptors.

These findings point to new therapeutic opportunities. Beyond immunosuppression, which induces remission in only ∼30–40% of patients^54^, our results highlight the potential of targeting H_2_ accumulation and *asr*-positive, multi-hydrogenase fermenters. This supports investigating dietary strategies^55^ that reduce fermentable substrates and thereby lower both H_2_ and H_2_S production, consistent with European Crohn’s and Colitis Organisation guidance on exclusive enteral nutrition^56^ and with reduced breath H_2_ in healthy individuals on low-fermentable-carbohydrate diets^57^.

Defining the sulfur sources used by these organisms, including dietary cysteine^22,33^, taurine from bile acids, mucosal sulfopolysaccharides, and other intestinal sulfite pools^13,30,32^, will be important for developing microbiome-informed interventions and biomarkers of disrupted intestinal H_2_ homeostasis. These findings provide a framework for developing microbiome-informed strategies to limit excess gas accumulation and H_2_S-mediated mucosal damage in Crohn’s disease.

## SUPPLEMENTAL FIGURES

**Fig. S1:**
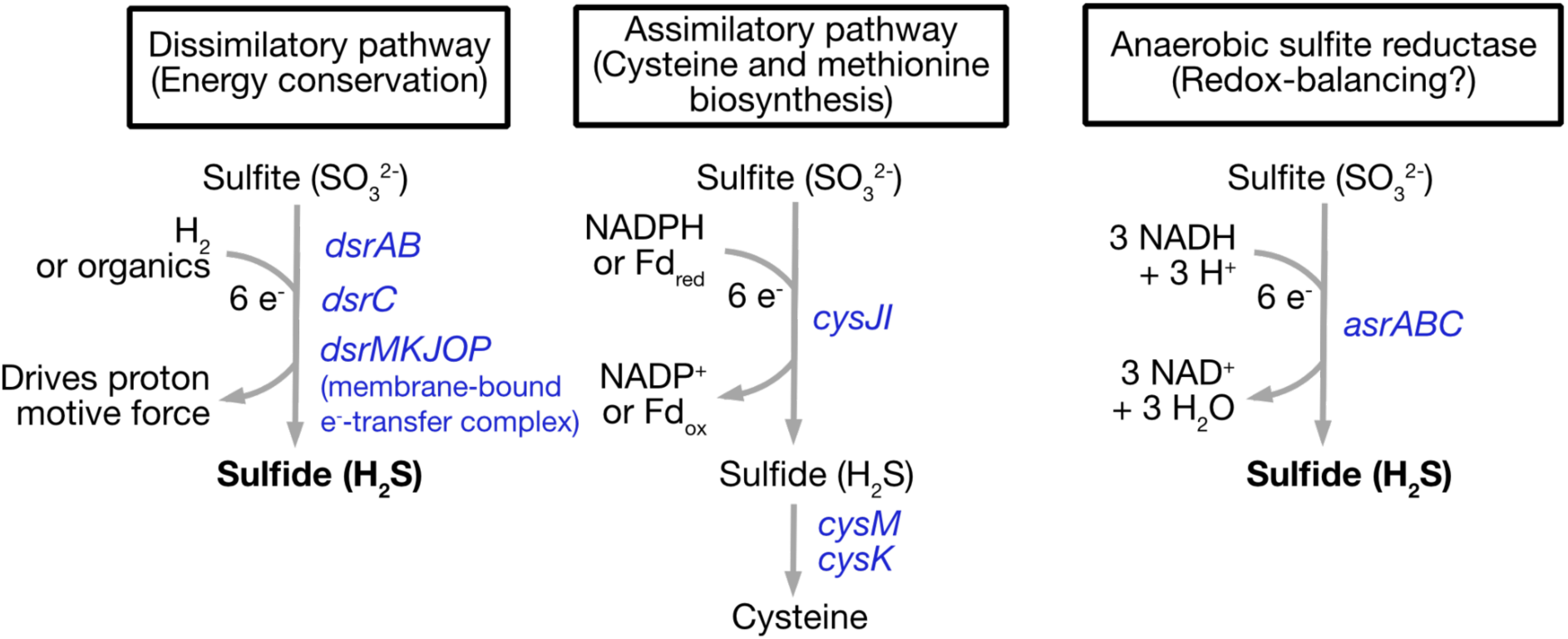
Sulfate reduction pathways. The dissimilatory (left) and assimilatory (middle) sulfate reduction pathway and Asr-mediated sulfate reduction (right) are shown, based on previous reports^58–60^. Microbial H_2_S production has been classically attributed to dissimilatory sulfate reduction (DSR) **(left)**. This dissimilatory pathway is the central energy-conserving step in the metabolism of classical Sulfate-Reducing Bacteria (SRB) such as *Desulfovibrio* and *Bilophila* species, which convert inorganic sulfate or sulfite to H_2_S. However, anaerobic sulfite reductases (Asr) also convert sulfite to H_2_S **(right)**. They use inorganic or organic substrates like cysteine, methionine, and taurine^31,32^ and have been suggested as a source of H_2_S formation in the gut^33,32,23,28^. Asr genes show resemblance to the sulfite reductase of the assimilatory sulfate reduction pathway for cysteine and methionine biosynthesis **(middle)**, and is sometimes considered part of this pathway^59^. However, *asr* is genetically distinct from its biosynthetic analog, uses NADH (generated in catabolic processes like fermentation) rather than NADPH as electron donor^49^, and is regulated by available electron acceptors (i.e., NAD^+^) rather than the cell’s cysteine demand^31,49^. Therefore, Asr may serve redox balancing by internally regenerating NAD^+^ from NADH, which is generated in catabolic processes such as fermentation.

**Fig. S2.**
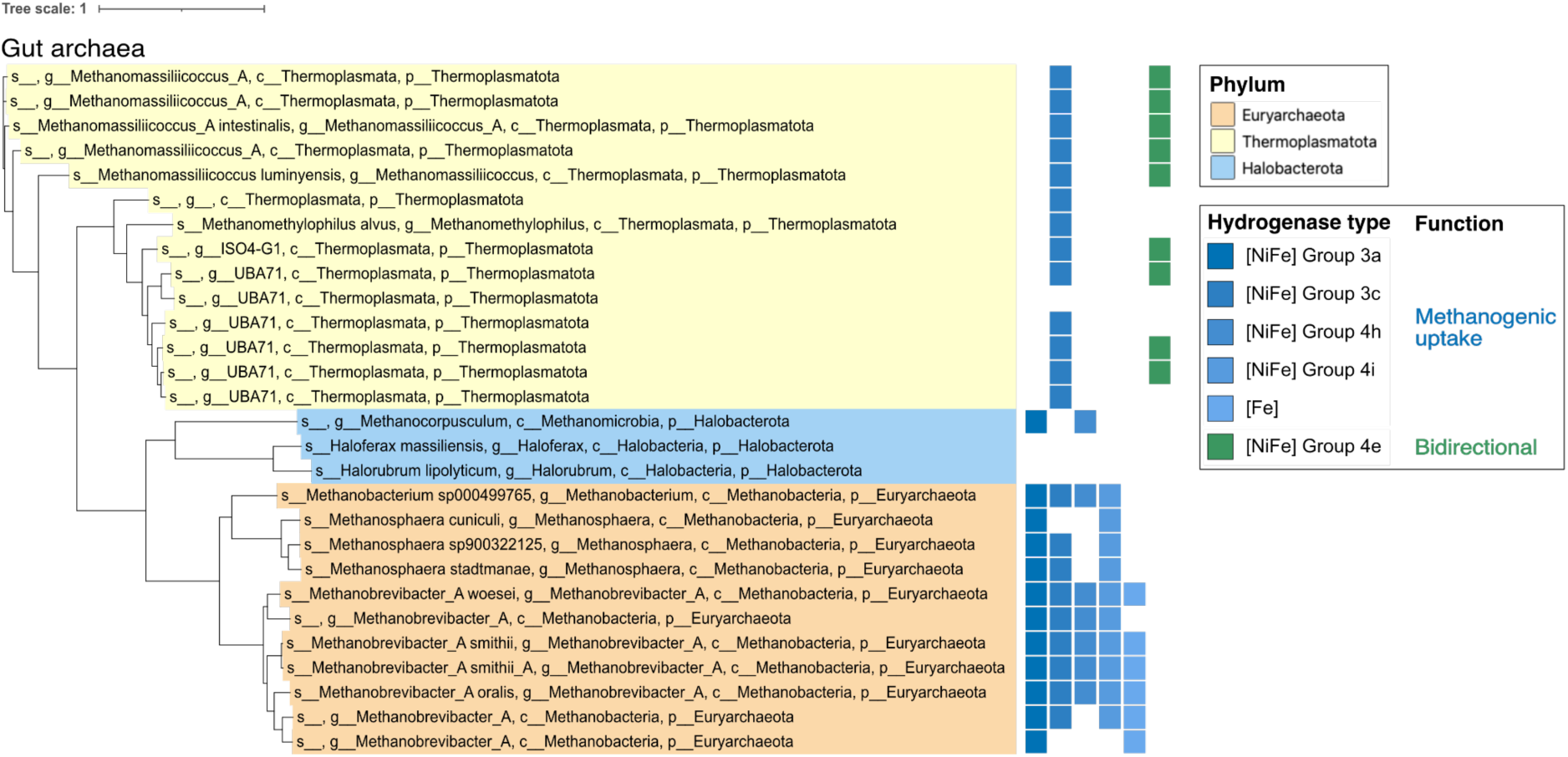
Phylogenetic distribution of hydrogenase types and associated function in archaea of the Unified Human Gastrointestinal Genome Collection (UHGG). Hydrogenases of methanogenic uptake, sometimes accompanied by bidirectional hydrogenases, were found.

**Fig. S3.**
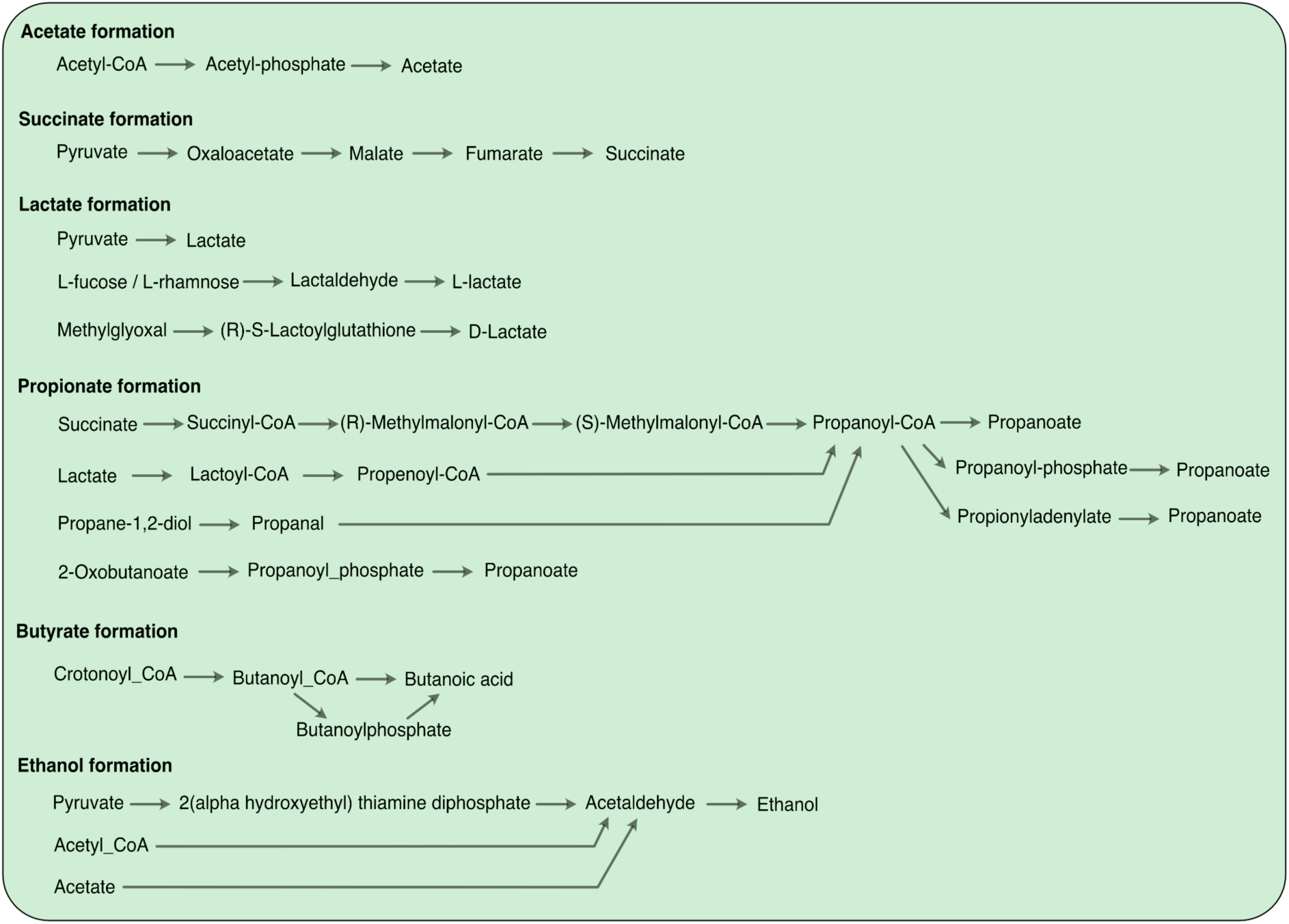
Fermentation pathways linked to hydrogen production. These fermentation pathways generate reduced electron carriers such as NADH or reduced ferredoxin, which can be oxidised by hydrogenases to reduce protons, releasing hydrogen gas. Based on KEGG Pathway Maps^61^, MetaCyc Pathways^62^, and Hackmann *et al.*^21^.

**Fig. S4.**
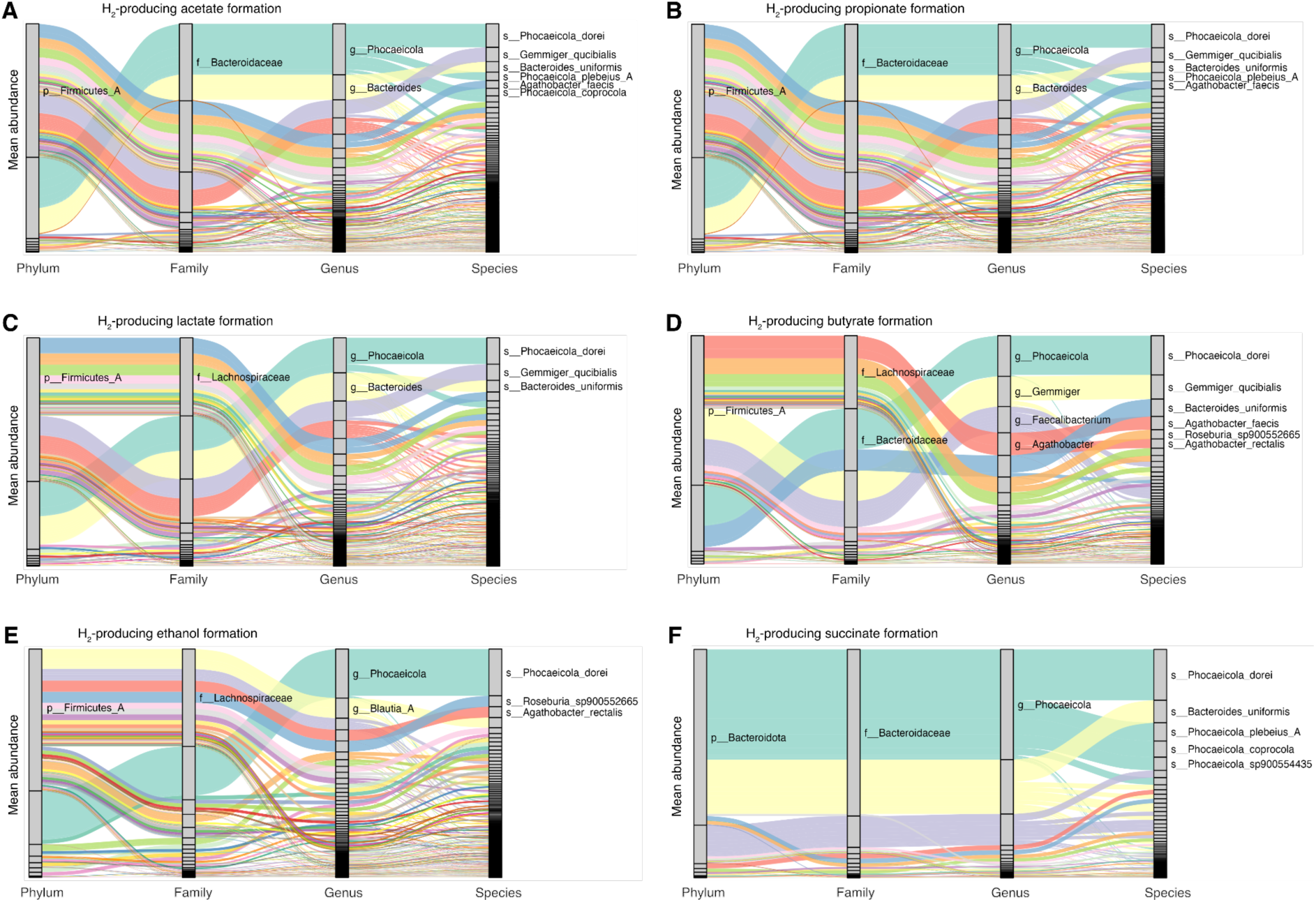
Relative abundance of H_2_ producers associated with each H_2_-producing pathway. Mean abundance and taxonomic identity of the H_2_ producers capable of (A) acetate formation, (B) propionate formation, (C) lactate formation, (D) butyrate formation, (E) ethanol formation, and (F) succinate formation.

**Fig. S5.**
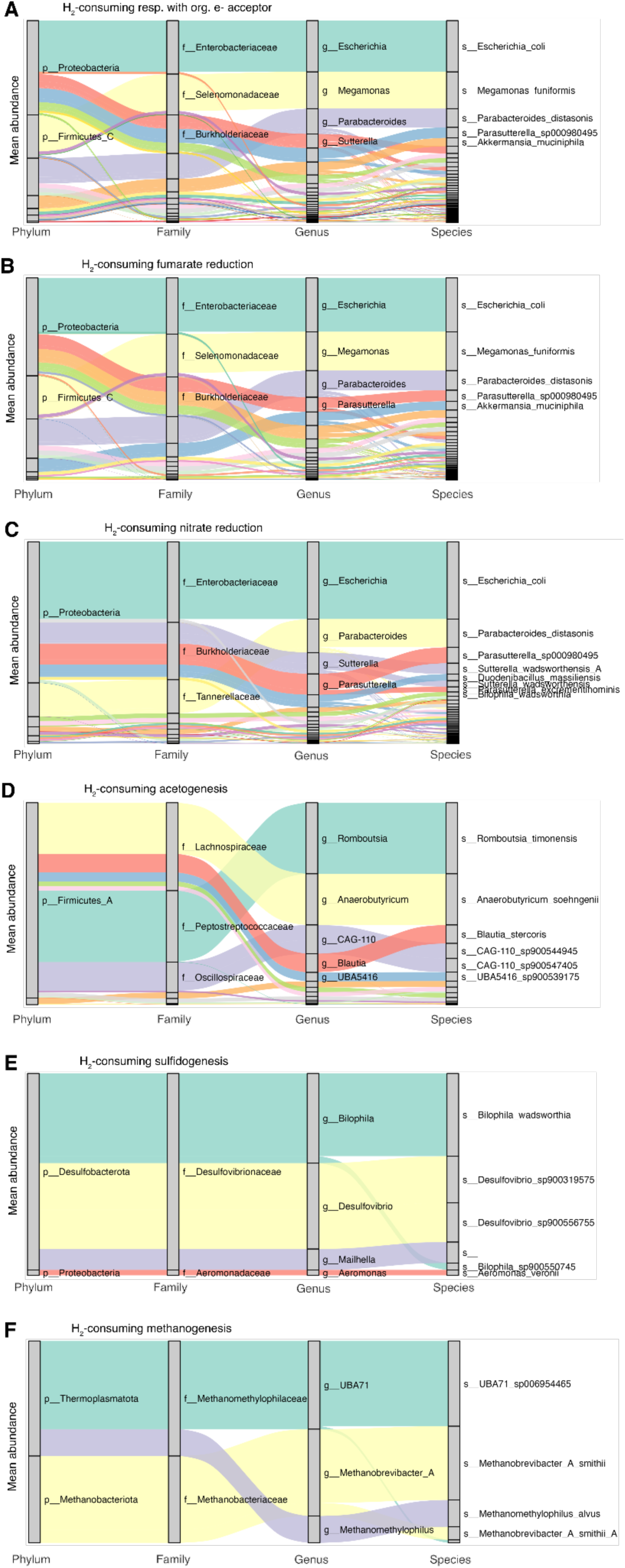
Relative abundance of H_2_ consumers and flexibles associated with each H_2_-consuming pathway. Mean abundance and taxonomic identity of the H_2_ consumers and flexibles capable of (A) anaerobic respiration with organic electron acceptors, (B) specifically fumarate reduction, (C) (partial) denitrification, (D) acetogenesis, (E) sulfidogenesis, and (F) methanogenesis.

**Fig. S6.**
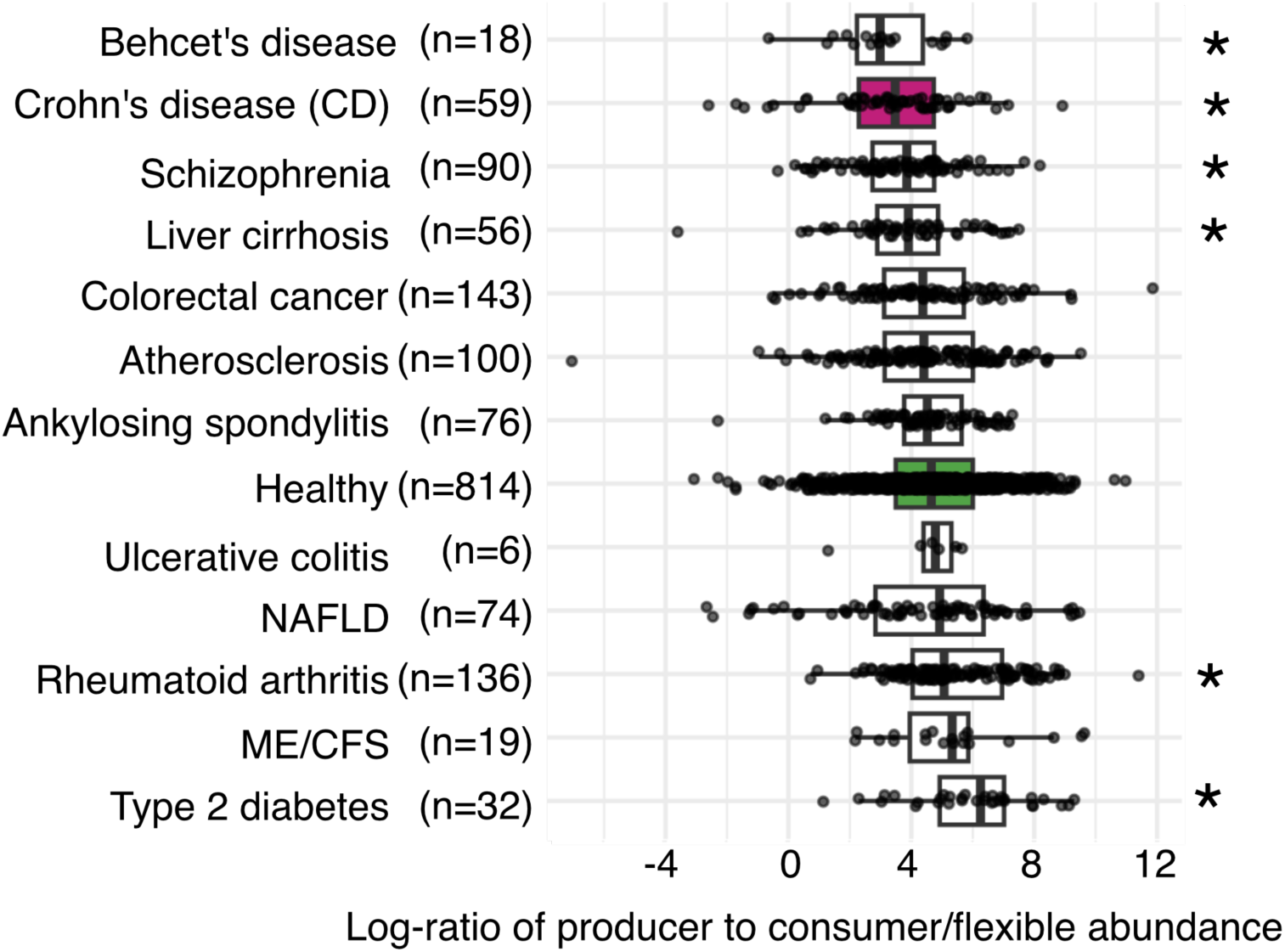
Log2-ratio of H_2_ producer abundance per H_2_ consumer and flexible abundance for healthy and diseased cohorts. Points show the log2-ratio for each individual, with n as the number of individuals per cohort, and box plots indicate the median and interquartile range. Boxplots were sorted by median. Statistical differences between groups were assessed using a Wilcoxon rank-sum test, where a star represents *p* < 0.05 (*p*-values 4×10⁻⁶ (schizophrenia), 10^−5^ (Crohn’s disease), 4×10⁻⁴ (rheumatoid arthritis), 5×10⁻⁴ (type2 diabetes), 7×10⁻⁴ (Behcet’s disease), 6×10⁻³ (liver cirrhosis), 0.2 (atherosclerosis), 0.2 (colorectal cancer), 0.3 (ME/CFS), 0.7 (ankylosing spondylitis), 0.7 (NAFLD), and 0.9 (ulcerative colitis)).

**Fig. S7.**
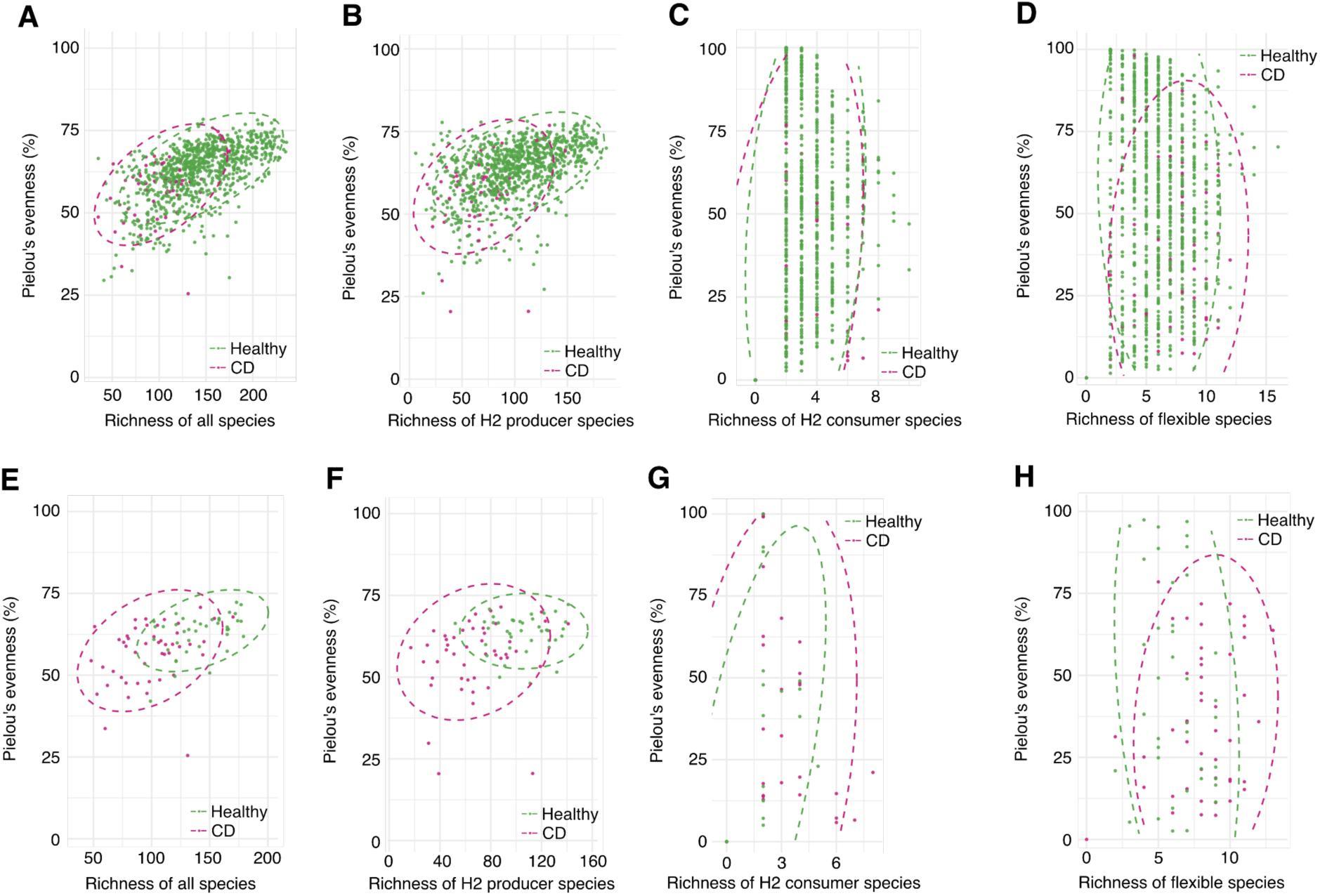
Species richness (x-axis) and Pielou’s evenness index (y-axis) for Crohn’s disease patients and healthy controls. Stool samples from all studies (A-D) and from the controlled He. et al study (E-H) are shown, considering all species, H_2_ producer species, H_2_ consumer species, and flexible species.

**Fig. S8.**
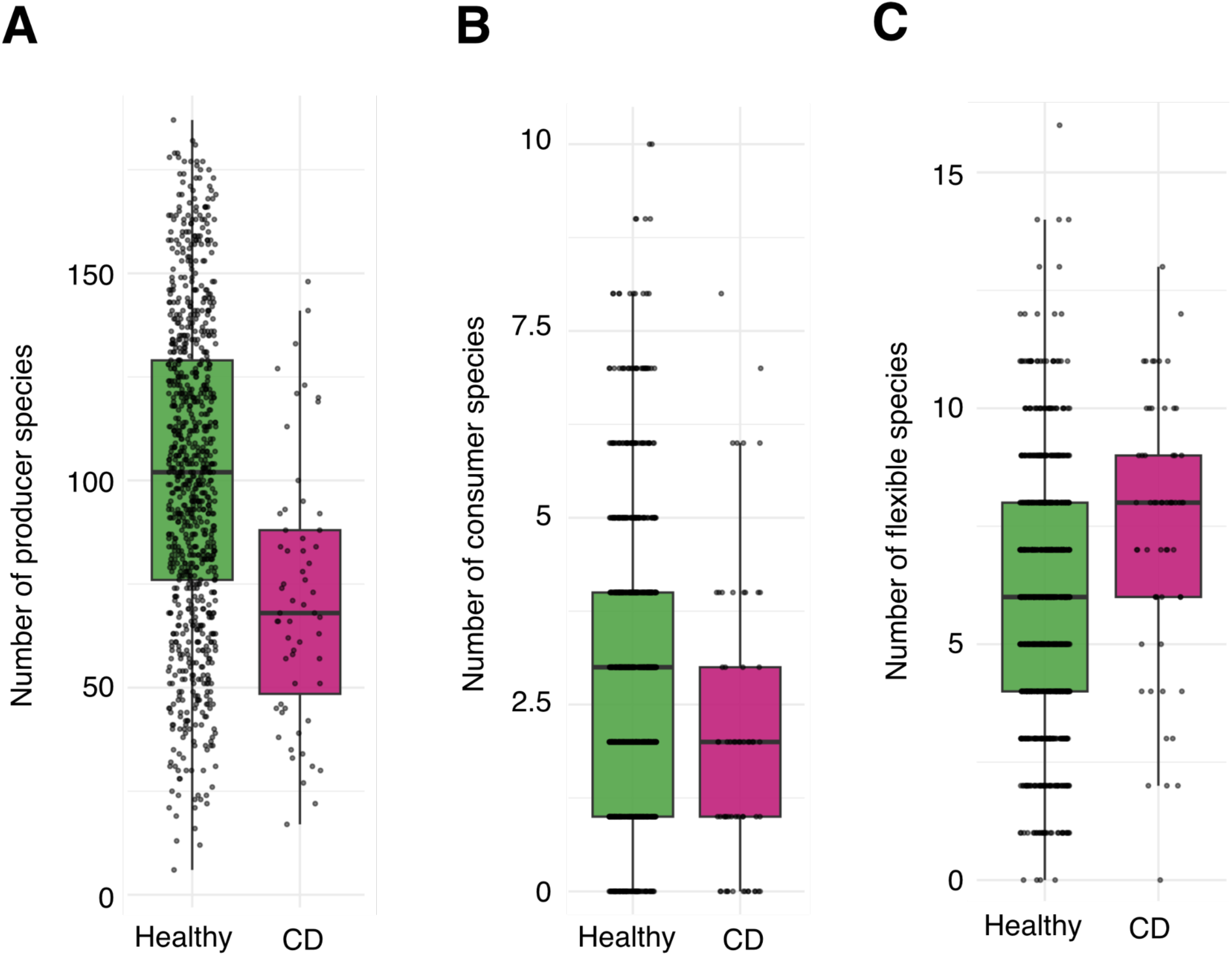
Richness of H_2_ producers and H_2_ consumers in healthy individuals and Crohn’s disease patients. The richness is the number of (A) H_2_ producer species, (B) H_2_ consumer species, and (C) flexible species that were detected in each patient’s stool sample.

**Fig. S9.**
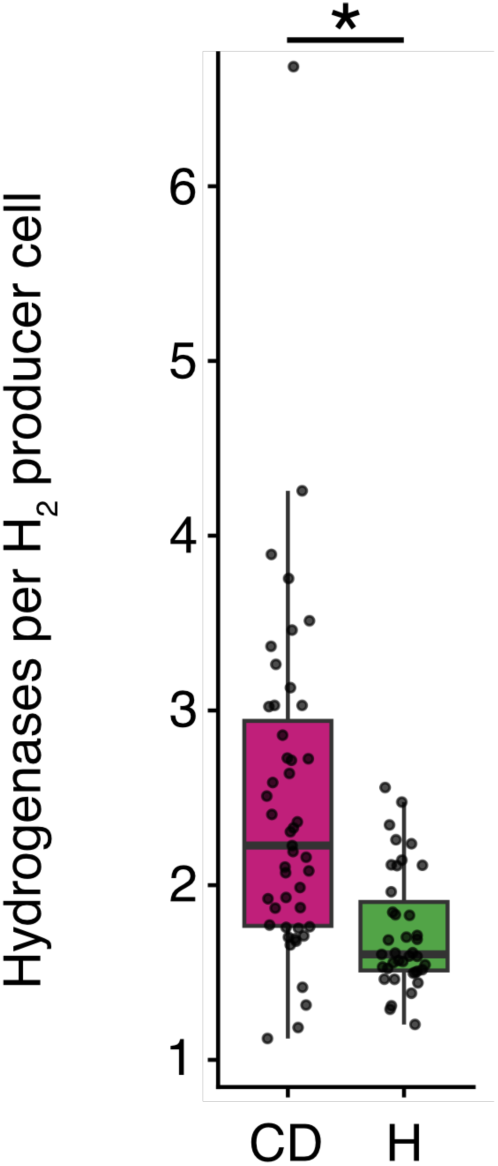
Average number of hydrogenases per genome in H_2_ producers in Crohn’s disease (CD) patients and healthy individuals (H). Each dot is the number of hydrogenase genes per genome, normalized by the abundance of each H_2_ producer genome (n healthy = 39, n CD = 46; Wilcoxon rank-sum test, *p* = 3×10^−6^).

**Fig. S10.**
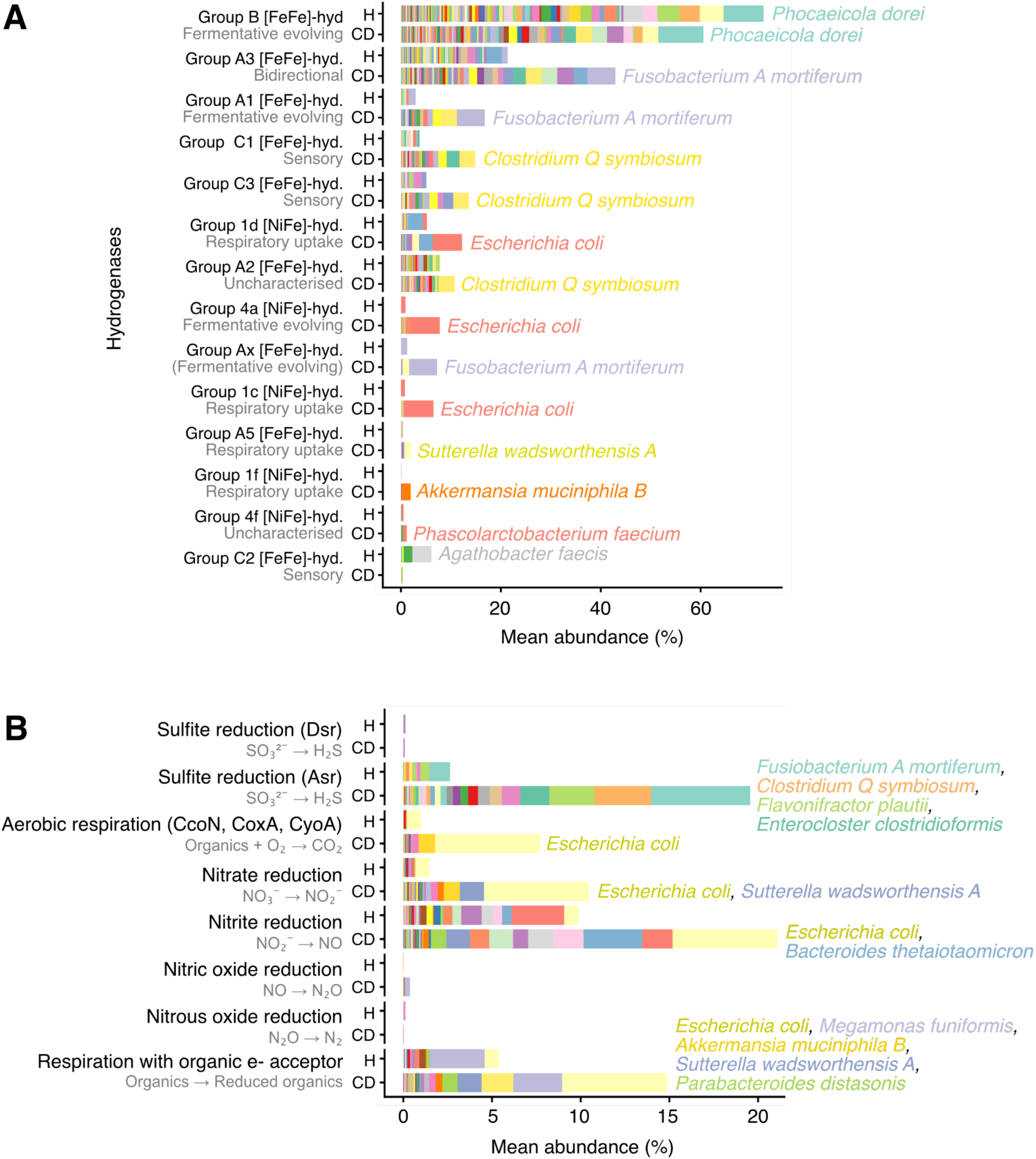
Relative abundances of taxa possessing hydrogenases (A) and genes connected to H_2_S production, aerobic respiration, denitrification, and DNRA (B). Plots correspond to Figure 4G and I, with the most abundant species annotated in the matching colour with the bar plot. Taxonomic names of the most abundant species in CD are displayed in corresponding colours to the bar plot. H = healthy, CD = Crohn’s disease.

**Fig. S11.**
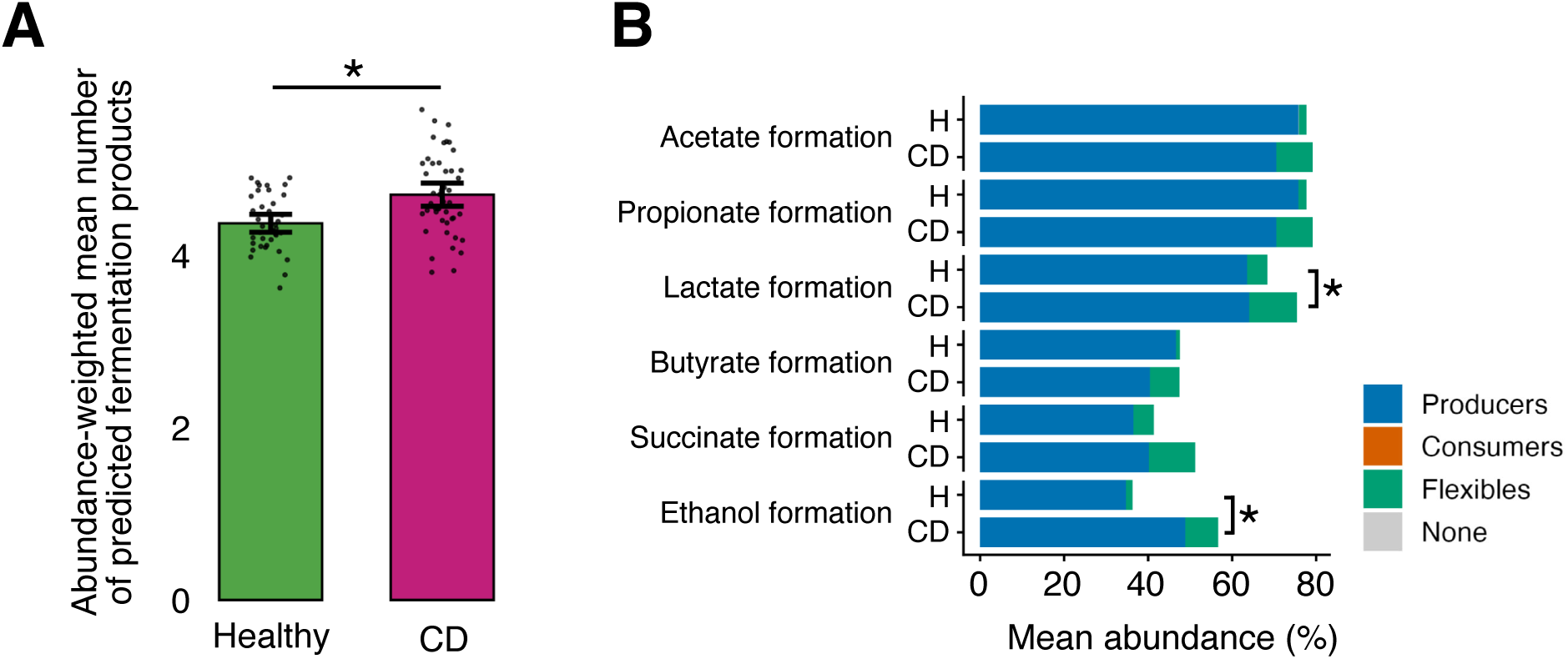
Increased diversity of fermentation capabilities in CD and shift towards ethanol and lactate formers. (A) Average number of predicted fermentation products in H_2_ producers, weighted by their respective abundance in CD patients and healthy controls (n_Healthy_ = 39, n_CD_ = 47; Wilcoxon rank-sum test, *p* = 5×10^−4^. (B) Abundance of taxa with fermentation capacities and H_2_-producing hydrogenases in stool sample from CD patients and healthy controls (n_Healthy_ = 39, n_CD_ = 46; Wilcoxon rank-sum test, *p*-values 0.3 (acetate formation), 0.3 (propionate formation), 0.03 (lactate formation), 0.9 (butyrate formation), 0.07 (succinate formation), 3×10^−5^ (ethanol formation)). H = healthy, CD = Crohn’s disease.

## SUPPLEMENTAL TABLES

**Table S1.** Hydrogenase types and their function.

| Hydrogenase type | Function | Reference |
| --- | --- | --- |
| [Fe] | Methanogenic uptake | 63 |
| Group A1 [FeFe] | Fermentative evolving | 35,44 |
| Group A2 [FeFe] | Unknown | 35,44 |
| Group A3 [FeFe] | Bidirectional | 35,44 |
| Group A4 [FeFe] | Bidirectional | 35,44 |
| Group A5 [FeFe] | Respiratory uptake | 64,65 , this study |
| Group A6 [FeFe] | Acetogenic uptake | 66 , this study |
| Group B [FeFe] | Fermentative evolving | 12 |
| Group C1 [FeFe] | Sensory (putative) | 35,38 |
| Group C2 [FeFe] | Sensory (putative) | 35,38 |
| Group C3 [FeFe] | Sensory (putative) | 35,38 |
| Group 1a [NiFe] | Respiratory uptake | 32 |
| Group 1b [NiFe] | Respiratory uptake | 32 |
| Group 1c [NiFe] | Respiratory uptake | 32 |
| Group 1d [NiFe] | Respiratory uptake | 32 |
| Group 1e [NiFe] | Respiratory uptake | 32 |
| Group 1f [NiFe] | Respiratory uptake | 32 |
| Group 1g [NiFe] | Respiratory uptake | 32 |
| Group 1h [NiFe] | Respiratory uptake | 35,67 |
| Group 1i [NiFe] | Respiratory uptake | 38 |
| Group 1j [NiFe] | Respiratory uptake | 38 |
| Group 1k [NiFe] | Methanogenic uptake | 38 |
| Group 1l [NiFe] | Respiratory uptake | 68 |
| Group 2a [NiFe] | Respiratory uptake | 63 |
| Group 2b [NiFe] | Sensory | 63 |
| Group 2c [NiFe] | Sensory | 32 |
| Group 2d [NiFe] | Unknown | 32 |
| Group 2e [NiFe] | Respiratory uptake | 38 |
| Group 3a [NiFe] | Methanogenic uptake | 63 |
| Group 3b [NiFe] | Bidirectional | 63 |
| Group 3c [NiFe] | Methanogenic uptake | 63 |
| Group 3d [NiFe] | Bidirectional | 63 |
| Group 4a [NiFe] | Fermentative evolving | 32 |
| Group 4b [NiFe] | Respiratory evolving | 35,38 |
| Group 4c [NiFe] | Respiratory evolving | 32 |
| Group 4d [NiFe] | Respiratory evolving | 35,38 |
| Group 4e [NiFe] | Bidirectional | 32 |
| Group 4f [NiFe] | Unknown | 32 |
| Group 4g [NiFe] | Unknown | 38 |
| Group 4h [NiFe] | Methanogenic uptake | 38 |
| Group 4i [NiFe] | Methanogenic uptake | 38 |

**Table S2. Functional annotation and classification of the Unified Human Gastrointestinal Genome (UHGG) collection.** Genomes listed alongside their taxonomy, predicted metabolic capabilities, and classification as H_2_ producers, H_2_ consumers, H_2_ flexibles, or non-H_2_-cyclers. *(In Supplemental Data)*

**Table S3.** Presence of H_2_ producing and consuming capabilities per phylum. Number of species states the number of UHGG species per phylum. Regarding Bacteroidota, there was a wide diversity of non-H_2_-metabolising Bacteroidota species, though the H2-metabolising species have been found to be more abundant in the human gut^12^. Archaeal phyla in italic.

| Phylum | Species (n) | Producer (%) | Consumer (%) | Flexible (%) | None (%) |
| --- | --- | --- | --- | --- | --- |
| Firmicutes_A | 1870 | 82.3 | 0.3 | 2.6 | 14.9 |
| Actinobacteriota | 828 | 71 | 1.9 | 9.7 | 17.4 |
| Bacteroidota | 603 | 23.9 | 2 | 3.3 | 70.8 |
| Firmicutes | 488 | 18.4 | 1.6 | 0.2 | 79.7 |
| Proteobacteria | 349 | 24.1 | 13.5 | 26.6 | 35.8 |
| Firmicutes_C | 148 | 50 | 4.7 | 27.7 | 17.6 |
| Verrucomicrobiota | 66 | 31.8 | 13.6 | 1.5 | 53 |
| Cyanobacteria | 63 | 0 | 0 | 0 | 100 |
| Campylobacterota | 47 | 0 | 51.1 | 42.6 | 6.4 |
| Fusobacteriota | 35 | 60 | 0 | 0 | 40 |
| Desulfobacterota_A | 27 | 0 | 18.5 | 81.5 | 0 |
| Firmicutes_I | 25 | 24 | 16 | 0 | 60 |
| Spirochaetota | 19 | 73.7 | 0 | 0 | 26.3 |
| Firmicutes_B | 14 | 85.7 | 0 | 7.1 | 7.1 |
| *Thermoplasmatota* | 14 | 0 | 35.7 | 50 | 14.3 |
| *Euryarchaeota* | 11 | 0 | 81.8 | 0 | 18.2 |
| Patescibacteria | 11 | 0 | 0 | 0 | 100 |
| Synergistota | 9 | 55.6 | 0 | 0 | 44.4 |
| Elusimicrobiota | 6 | 16.7 | 0 | 0 | 83.3 |
| Eremiobacterota | 4 | 75 | 0 | 0 | 25 |
| *Halobacterota* | 3 | 0 | 0 | 0 | 100 |
| Bdellovibrionota | 1 | 0 | 0 | 0 | 100 |
| Fibrobacterota | 1 | 0 | 0 | 0 | 100 |
| Firmicutes_G | 1 | 100 | 0 | 0 | 0 |
| Myxococcota | 1 | 100 | 0 | 0 | 0 |

**Table S4.**
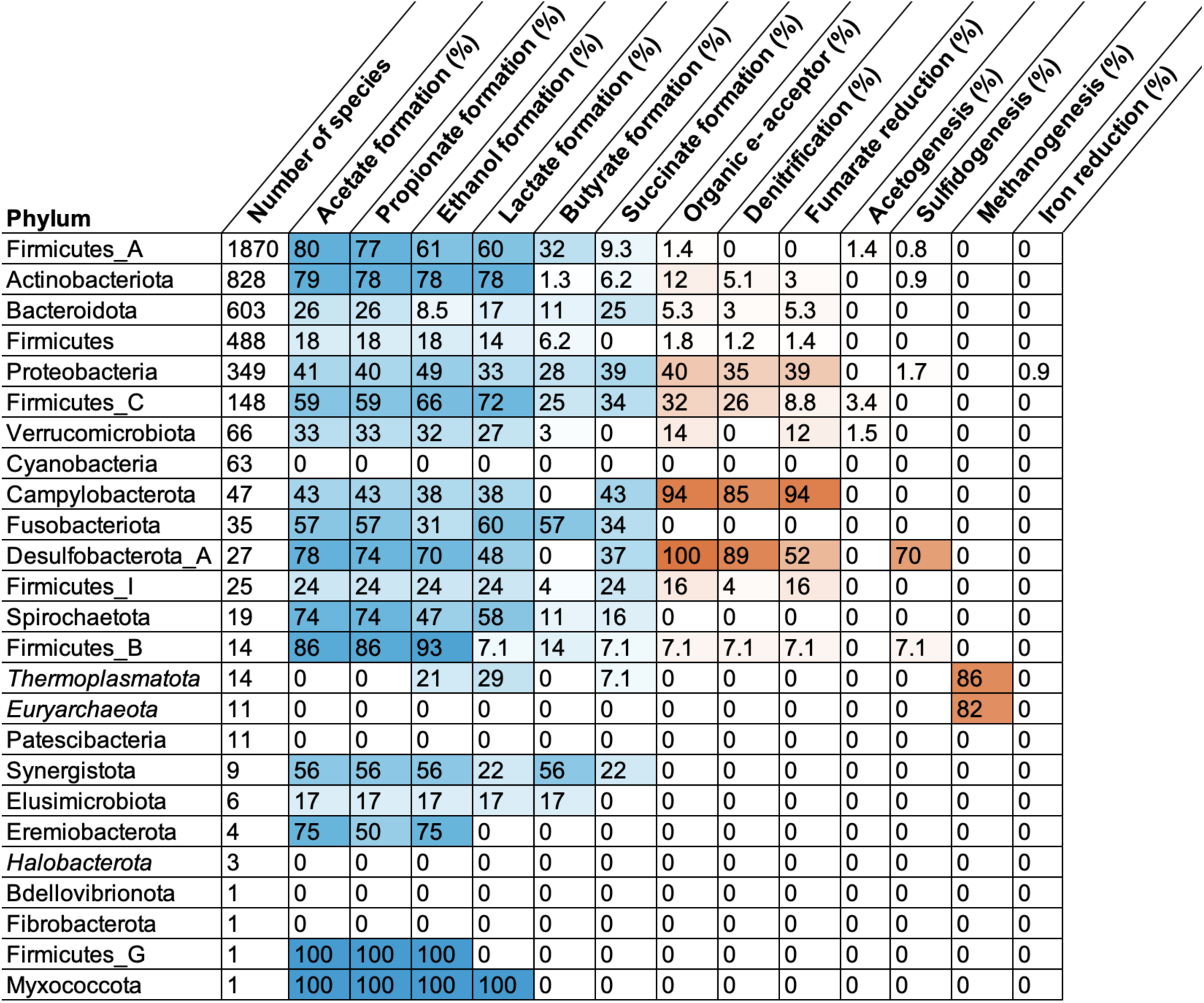
Presence of H_2_ producing and consuming capabilities per phylum. H_2_ producing and consuming capabilities are defined as the presence of H_2_-producing or -consuming hydrogenases in combination with potentially H_2_-producing or -consuming pathways, respectively. Number of species states the number of UHGG species per phylum. The use of organic electron acceptors including fumarate for respiration was common in the phyla Proteobacteria, Campylobacterota, and Desulfobacterota A. Methanogenesis was prevalently found in the phyla Thermoplasmatota and Euryarchaeota. Potential H_2_ consumption through sulfidogenesis was found to be most common in Desulfobacterota A species (in ca. 70% of species), Acetogenesis most common in Firmicutes C (in ca. 3% of species), and iron reduction only in Proteobacteria (in ca. 1% of species). Archaeal phyla in italic.

**Table S5. Functional annotation, H_2_-cycling capabilities, and relative abundances of the ca. 1,700 stool metagenomes.** Genomes listed alongside their taxonomy, predicted metabolic capabilities, classification as H_2_ producers, H_2_ consumers, H_2_ flexibles, or non-H_2_-cyclers, and abundances in each sample. Species-level MAG abundances per sample were estimated with KMA^69^ by Marcelino *et al.*^24^*. (In Supplemental Data)*

**Table S6. Abundance statistics and raw abundances of predicted H_2_ producers in the stool microbiota of 888 healthy individuals.** *(In Supplemental Data)*

**Table S7. Abundance statistics and raw abundances of predicted H_2_ consumers in the stool microbiota of 888 healthy individuals.** *(In Supplemental Data)*

**Table S8. Abundance statistics and raw abundances of predicted H_2_ flexibles in the stool microbiota of 888 healthy individuals.** *(In Supplemental Data)*

**Table S9.**
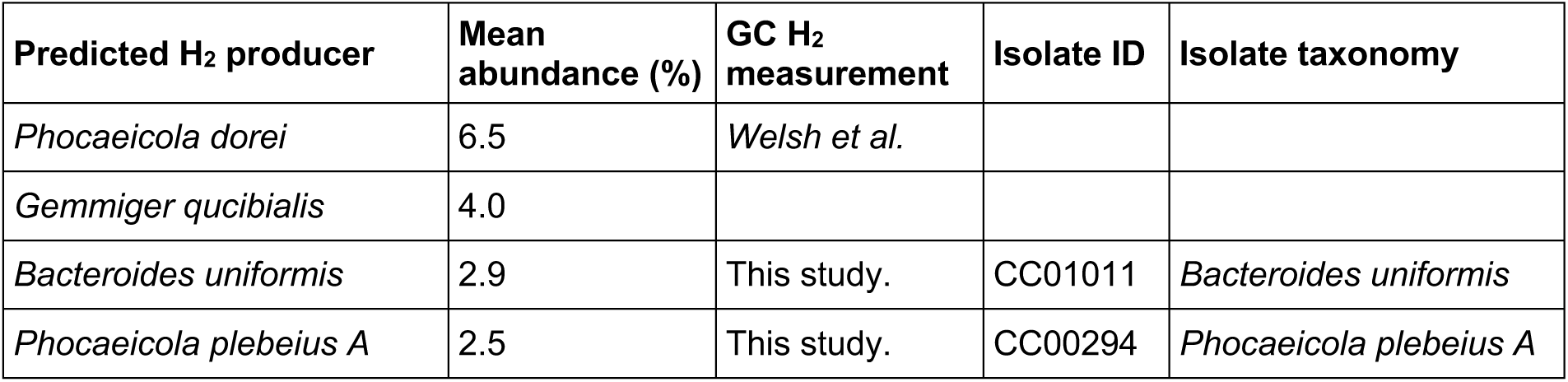
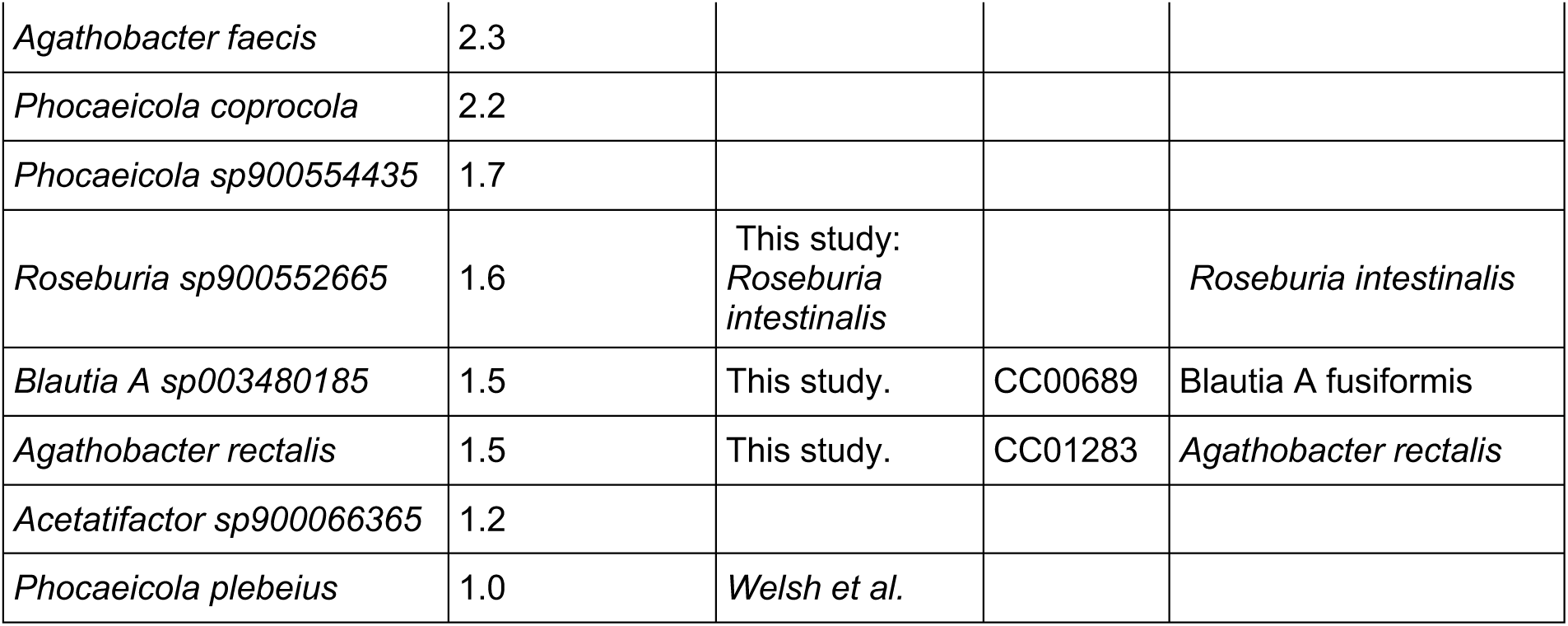
Measurements of H_2_ production of abundant gut bacteria. The H_2_ producer species with the highest mean abundance in the stool of 888 healthy individuals are shown, alongside their abundance, measurements of H_2_ production in this study or by Welsh *et al.*^12^, and reference to the here studied isolate.

**Table S10. Functional annotation, H_2_-cycling capabilities, and relative abundances of the genomes from stool metagenomes of healthy controls in the study of He *et al.***^70^. Genomes listed alongside their taxonomy, hydrogenases, predicted metabolic capabilities, classification as H_2_ producers, H_2_ consumers, H_2_ flexibles, or non-H_2_-cyclers, and abundances in each sample. *(In Supplemental Data)*

**Table S11. Functional annotation, H_2_-cycling capabilities, and relative abundances of the genomes from stool metagenomes of CD patients in the study of He *et al.***^70^. Genomes listed alongside their taxonomy, hydrogenases, predicted metabolic capabilities, classification as H_2_ producers, H_2_ consumers, H_2_ flexibles, or non-H_2_-cyclers, and abundances in each sample. *(In Supplemental Data)*

**Table S12. Significantly more abundant H_2_ producers in the stool metagenomes of CD patients compared to healthy controls in the study of He *et al.***^70^. Genomes are listed alongside their taxonomy and significance metrices. *(In Supplemental Data)*

**Table S13. Significantly less abundant H_2_ producers in the stool metagenomes of CD patients compared to healthy controls in the study of He *et al.***^70^. Genomes are listed alongside their taxonomy and significance metrices. *(In Supplemental Data)*

**Table S14. Significantly more abundant H_2_ consumers in the stool metagenomes of CD patients compared to healthy controls in the study of He *et al.***^70^. Genomes are listed alongside their taxonomy and significance metrices. *(In Supplemental Data)*

**Table S15. Significantly more abundant H_2_ flexibles in the stool metagenomes of CD patients compared to healthy controls in the study of He *et al.***^70^. Genomes are listed alongside their taxonomy and significance metrices. *(In Supplemental Data)*

**Table S16.** Measurements of H_2_ production during growth of the species with most increased abundance in CD patients. The H_2_ producer species with the highest mean abundance are shown, alongside their abundance, measurements of H_2_ production in this study or by Welsh *et al.*^12^, and reference to the here studied isolate.

| Predicted H<sub>2</sub> producer | Mean abundance (%) | GC H<sub>2</sub> measurement | Isolate ID | Isolate taxonomy |
| --- | --- | --- | --- | --- |
| *Clostridium Q symbiosum* |  | This study. | CC00921 | *Clostridium symbiosum* (*Otoolea symbiosa*) |
| *Enterocloster clostridioformis* |  | This study. | CC01278 | *Enterocloster clostridioformis* |
| *Erysipelatoclostridium ramosum* |  |  |  |  |
| *Hungatella effluvii* |  | This study. | CC00403 | *Hungatella effluvii* |
| *Fusobacterium A ulcerans* |  |  |  |  |
| *Clostridium AQ innocuum* |  | This study. | CC00281 | *Clostridium AQ innocuum* |
| *UBA9502 sp900540335* |  |  |  |  |
| *CAG-81 sp900066785* |  |  |  |  |
| *Bacteroides faecis* |  | Welsh *et al.* |  |  |
| *UBA1691 sp900544375* |  |  |  |  |

**Table S17.** H_2_-producing and -sensing hydrogenases in the stool-derived isolates. Corresponds to isolates tested for H2 production in Figure 3 and 4.

| Isolate name | Prevalent | FeFe Group A1 | FeFe Group A2 | FeFe Group A3 | FeFe Group B | FeFe Group C1 | FeFe Group C3 |
| --- | --- | --- | --- | --- | --- | --- | --- |
| *Agathobacter rectalis* | Healthy | 0 | 0 | 0 | 2 | 0 | 0 |
| *Bacteroides uniformis* | Healthy | 0 | 0 | 0 | 1 | 0 | 0 |
| *Blautia A sp003480185* | Healthy | 1 | 0 | 0 | 1 | 0 | 0 |
| *Phocaeicola plebeius A* | Healthy | 0 | 0 | 0 | 1 | 0 | 0 |
| *Clostridium AQ innocuum* | CD | 0 | 1 | 0 | 2 | 0 | 0 |
| *Enterocloster clostridioformis* | CD | 1 | 0 | 1 | 2 | 1 | 1 |
| *Hungatella effluvii* | CD | 1 | 0 | 2 | 1 | 1 | 1 |
| *Clostridium Q symbiosum* | CD | 1 | 0 | 1 | 1 | 1 | 2 |

**Table S18.** Biopsy sample information.

| Sample ID | Sex | Diagnosis | Details |
| --- | --- | --- | --- |
| B01 | M | Non_CD | IBS, non-inflamed |
| B02 | F | Non_CD | IBS, non-inflamed,<br>post-infectious IBS |
| B04 | F | CD | Newly diagnosed<br>Crohn's |
| B05 | F | Non_CD | Non-inflamed, PR<br>bleeding, worms in<br>colon |
| B06 | F | CD | Newly diagnosed<br>Crohn's |

## SUPPLEMENTARY NOTES

### Supplementary Note 1: Species richness and Pielou’s evenness index

The shift in the gut microbiota of CD patients has been associated with reduced microbial diversity. To test whether the diversity of H_2_ producers and consumers differed between healthy and CD patients, we computed their species richness and evenness. The species richness was measured as the number of species per stool sample, and the Pielou’s evenness index was calculated, where 1 indicated equal abundance of all species and values close to 0 indicated the dominance of few species over others.

Across all studies, the mean richness of CD patients was reduced from 144 species +/- 40 (mean +/- SD) in healthy individuals to 103 species +/- 33 (**Fig. S5A**). The Pielou’s evenness index was reduced from 0.63 +/- 0.08 in healthy to 0.58 +/- 0.09 in CD patients. The species richness of the H_2_ producers specifically shrank from 102 +/- 37 in healthy to 72 +/- 31 in CD patients (**Fig. 4B**), and the richness of H_2_ consumers and flexibles remained similar (2.8 +/- 1.9 to 2.1 +/- 2.0 in H_2_ consumers, 5.9 +/- 2.5 to 7.5 +/- 2.8 in flexibles) (**Fig. S5C and S5D**). The Pielou’s evenness index of both H_2_ producers and H_2_ consumers and flexibles sank slightly (for H_2_ producers from 0.63 +/- 0.08 in healthy to 0.57 +/- 0.11 in CD patients; for H_2_ consumers and flexibles from 0.54 +/- 0.22 in healthy to 0.44 +/- 0.22 in CD patients) (**Fig. 4B, S5**).

When focusing only one study of CD patients and healthy case-controls (He et al. study) to remove confounding factors such as ethnicity and study design, the mean richness in CD patients’ stool samples was reduced from 142 species +/- 25 (mean +/- SD) in healthy individuals to 99 species +/- 28. The Pielou’s evenness index was reduced from 0.63 +/- 0.06 in healthy to 0.57 +/- 0.09 in CD patients (**Fig. S5E**). The species richness of H_2_ producers specifically shrank from 106 +/- 22 in healthy to 68 +/- 28 in CD patients (**Fig. S5F**), while the species richness of H_2_ consumers and flexibles increased slightly (1.5 +/- 1.2 to 2.2 +/- 2.0 in consumers, 6.2 +/- 1.9 to 8.1 +/- 2.5 in flexibles) (**Fig. S5G and S5H**). The Pielou’s evenness index of both H_2_ producers, and H_2_ consumers and flexibles remained similar (for H_2_ producers 0.63 +/- 0.06 in healthy to 0.56 +/- 0.11 in CD patients, for H_2_ consumers and flexibles 0.43 +/- 0.24 in healthy and 0.42 +/- 0.20 in CD patients) (**Fig. 4B, S5**).

### Supplementary Note 2: Overproportional decrease in H_2_ producers in CD

To determine whether H_2_ cycling species were overproportionally affected by the abundance changes compared to non-H_2_-cycling species, we computed their enrichment within the species with increased or decreased abundance in CD patients. We found that H_2_ producers were overrepresented in the species with decreased abundance in CD patients (BH-adjusted *p* = 0.0001, NES = 1.7). Flexible species, that were predicted to be capable of both H_2_ production and consumption, were overrepresented in the species with increased abundance in CD patients (BH-adjusted *p* = 0.0005, NES = -2.2). Non-H_2_-cycling and H_2_ consuming species were not significantly overrepresented in the species with increased or decreased abundance in CD patients.

## Materials and Methods

### Gastrointestinal genome collection

As a comprehensive collection of human gut bacteria and archaea, we used the Unified Human Gastrointestinal Genome (UHGG) collection^37^. It is a comprehensive gut genome resource that combines 286,997 genomes representing 4,644 prokaryotic species. Taxonomy was assigned using GTDB-Tk v0.3.1^71^. The UHGG catalogue of 4,644 species representatives was downloaded from the ENA project PRJEB33885.

### Stool metagenomic datasets

To compare the microbiota of healthy and diseased patients, we analysed a stool metagenomic dataset that was assembled through a global survey of gut metagenomes from healthy and perturbed gut states^24^. It comprised 1,697 gut metagenome samples, including 888 healthy and 809 diseased individuals from 33 published studies, 15 countries and 11 disease phenotypes. It excluded studies with dietary interventions, medications, and exercise. It contains only one sample per individual and excludes samples of low sequencing depth (less than 15 M paired end reads after quality control). The healthy cohort included male and female individuals with a Body Mass Index (BMI) between 18.5 and 24.9 and no reported disease. Sequencing read assembly and binning had resulted in 24,369 high-quality MAGs (>90% completeness, <5% contamination, ENA PRJEB63093), which were dereplicated at 95% Average Nucleotide Identity (ANI) into 949 bacterial and 6 archaeal species representatives. These genomes were downloaded from Zenodo (https://zenodo.org/record/8223163; spp_level_representative_MAGs.zip). They were taxonomically classified using GTDB-Tk v.1.5.1^71^ and species-level MAG abundances per sample were estimated with KMA^69^ (**Tables 5**).

### Hydrogenase identification and classification

For identification of hydrogenase types in bacterial and archaeal genomes, we screened the predicted proteins using DIAMOND v0.9.22 (blastx) for alignment against the in-house curated hydrogenase database (github.com/GreeningLab/GreeningLab-database). Gene hits were filtered as by Lappan *et al.*^72^: Alignments were filtered for at least 80% query or 80% subject coverage, and then further filtered by a minimum percentage identity score of 60% for [FeFe]-hydrogenases and group 4 [NiFe]-hydrogenases, and 50% for [Fe]-hydrogenase and all other [NiFe]-hydrogenases. The types of group A [FeFe]-hydrogenases were previously reported to not be classifiable by sequence alone as they have principally diversified through changes in domain architecture and quaternary structure^44,35,38^. Therefore, the types of group A [FeFe]-hydrogenases were assigned based on the presence of conserved domains within the hydrogenase gene itself or the 2 genes upstream and downstream of the group A hydrogenase. Conserved domains were screened for with CD-search (rpsblast v2.15.0 with -evalue 1e-3, Conserved Domain Database (CDD) v3.21 was downloaded from ftp.ncbi.nlm.nih.gov/pub/mmdb/cdd/little_endian). Types of group A [FeFe]-hydrogenases were assigned based on Greening *et al.*^35^ as follows: Group A6 (acetogenic) [FeFe]-hydrogenase if the genome contained the genes AcsB or CooS (see next paragraph for their annotation); group A2 [FeFe]-hydrogenase if the conserved domains PRK12771, COG0493, PRK12814 (glutamate synthase domains) or PRK12831 (putative oxireductase domain) were detected; group A3 [FeFe]-hydrogenase if COG1894, TIGR01959, PRK13596, PRK11278 (NuoF subunit domains of NADH-quinone oxireductase) were detected; group A4 [FeFe]-hydrogenase if COG1142 (HycB), cd10554 (HycB-like), or PRK12769 (putative oxireductase Fe-S binding subunit) were detected; group A5 [FeFe]-hydrogenase if pfam02256, smart00902 (iron hydrogenase small subunit) were detected; group Ax (subtype unknown) [FeFe]-hydrogenase was annotated if flanking genes were missing in a genome due to incomplete sequencing; and all remaining group A [FeFe]-hydrogenases were assigned to be a group A1 (stand-alone) [FeFe]-hydrogenase.

### Visualisation of phylogenetic spread of hydrogenases

To visualise differences in the presence of hydrogenase types across bacterial and archaeal taxa, the phylogenetic tree of the UHGG collection from Almeida *et al.*^37^ was overlaid with the presence of hydrogenase types in each species-level cluster using the Interactive Tree of Life (iTOL)^73^ (v6, symbol height factor 20x for bacteria).

### Annotation of pathways linked to H_2_ production and H_2_ consumption

As potential H_2_-producing pathways, we considered fermentation pathways for acetate formation, propionate formation, ethanol formation, lactate formation, butyrate formation, and succinate formation, that may balance involved redox reactions through hydrogen formation. To detect pathways that are linked to H_2_ production, we annotated manually curated fermentation pathways that are linked to producing electrons and thus, potentially, H_2_ production. We used DRAM v1 (--min_contig_size 1000 --prodigal_mode meta)^74^ for functional annotation of KEGG’s KO numbers^61^, and filtered for DRAM hits of category *A (reciprocal best hit to a KEGG gene)*. We then searched for the presence of the genes in the manually curated pathways shown in Fig. S1.

Based on KEGG Pathway Maps^61^ (Last updated: December 8, 2025), MetaCyc Pathways^62^ (v29.5), and Hackmann et al.^21^, we searched for the following KO numbers of enzymes involved in fermentation and potentially linked to H_2_ production: acetate formation (K13788, K00625, K15024, K04020, K00925), butyrate formation (crotonoyl CoA to butyrate K00248, K00249, K06445, K09478, K00232, K09478, K01034, K01035, K19709; or K00248, K00249, K06445, K00232, K17829, K00209, K09478, K00634, K00929), succinate formation (K03416, K17489, K01676, K01677, K01678, K01679, K01675, K01774, K01675, K01774, K00244, K00245, K00246, K00247, K00239, K00240, K00241, K00247), actate formation (K00016, K03777, K03778, K03778, K03777; or K07248, K19266; or K01069, K05523), propionate formation (propanoyl-CoA formation K22214, K18118, K01899, K01900, K01902, K01903, K01899, K01900, K01847, K01849, K01848, K11942, K05606, K05606, K11264, K18426, K01604, K18426, K01905, K22224; or K01026, K20626, K20627, K00248, K00232, K19745, K20143; or K01699, K13919, K13920, K13922; propanoyl-CoA to propionate K01905, K22224, K24012, K01026, K13788, K00625, K15024, K04020, K13923, K00925, K00932, K19697, K01895, K01908; 2-oxobutanoate to propionate K00925, K00932, K19697), ethanol formation (K00132, K04021, K04072, K04073, K18366, K04072, K01568, K00128, K14085, K00149, K00138, K00001, K00121, K04072, K13951, K13952, K13953, K13954, K13980, K18857, K14028, K14029, K00002, K12957, K13979, K00114, K04022). We then considered a pathway to be present if at least half of the genes of the manually curated pathways, displayed in Fig. S3, were detected.

As potential H_2_-consuming pathways, we considered dissimilatory nitrate reduction to ammonium (DNRA), (partial) denitrification, fumarate reduction, acetogenesis, sulfidogenesis, methanogenesis, iron reduction, and anaerobic respiration with organic electron acceptors, such as TMSO, DMSO, and host-derived or dietary compounds as electron acceptors. To detect pathways that are linked to H_2_ consumption, we combined several annotation approaches. To identify respiration pathways with dietary or human-derived organic electron acceptors, molybdopterin and flavin reductases were identified by searching Pfam database HMMs (PF00384 and PF00890, respectively) with the tool Hmmsearch^75^ (v3.4; e-value cut-off 1e-5) based on Little *et al.*^10^. In addition, we screened for DMSO reduction by the presence of K07306 (*dmsA*) and for TMSO reduction by the presence of K07811 (*torA*), based on the above-described annotation of KO numbers through DRAM. To identify fumarate reduction specifically, denitrification, acetogenesis, sulfidogenesis, methanogenesis, and iron reduction, we screened the predicted proteins of all genomes using DIAMOND v0.9.22^76^ (blastx) for alignment against the in-house curated marker gene database (github.com/GreeningLab/GreeningLab-database). This included marker genes for fumarate reduction (*frdA*, which is however hard to distinguish from *sdhA* for succinate oxidation), reductive acetogenesis (*acsB*), dissimilatory sulfite reduction of sulfite (SO₃²⁻) to hydrogen sulfide (H_2_S) (*asrA*, dissimilatory sulfite reductase subunit A), anaerobic sulfite reduction to H_2_S (*asrA*, anaerobic sulfite reductase subunit A), nitrate reduction to nitrite (*narG, napA*), nitrite reduction to ammonium (*nrfA*), nitric oxide reduction to nitrous oxide (*norB*), nitrous oxide reduction to nitrogen (NosZ), methanogenesis (*mcrA*), and iron reduction (*mtrB, omcB*). We also screened for marker genes for aerobic respiration (*ccoN, coxA, cyoA*). Gene hits were filtered as by Lappan et al.^72^: Alignments were filtered for at least 80% query or 80% subject coverage, and then further filtered by a minimum percentage identity score by protein of 50%.

### Visualisation of combinations of H_2_-producing and consuming capabilities

As described above, a species was assigned to have H_2_ producing capabilities if its genome (i) harboured a H_2_-evolving hydrogenase AND (ii) harboured a potentially H_2_-producing fermentation pathway. A species was assigned to have H_2_ consuming capabilities if its genome (i) harboured a H_2_-uptaking hydrogenase AND (ii) harboured a potentially H_2_-consuming metabolic pathway. We considered bidirectional hydrogenases to enable H_2_-production in environment of the human gut, and did not include unclassified hydrogenases (group A2 [FeFe], 4f [NiFe], and 4g [NiFe] hydrogenases), thus potentially underestimating the number of H_2_ producers and consumers. The R package upSet complex^77,78^ (R v4.5.1) was used to visualise the combinations of H_2_-producing and H_2_-consuming capabilities of each species of the UHGG.

### Culturing of isolates

Isolates of gut bacteria were obtained from the Australian Microbiome Culture Collection (CC isolate IDs listed in Table S9 and S16). Isolates from glycerol stocks were plated on pre-reduced YCFA agar plates and grown anaerobically at 37°C until colonies formed. A single colony was used to inoculate a 5 ml of pre-reduced YCFA broth in a 15 mL Falcon tube, which was incubated anaerobically at 37°C. After overnight incubation, each pre-starter culture was transferred into fresh YCFA broth and grown to the log-phase. Log-phase cultures were then used to inoculate triplicate 30 mL aliquots of YCFA broth to a starting OD600 of 0.025. Cultures were maintained in 120 mL glass serum vials sealed with lab-grade butyl rubber stoppers for hydrogen production measurements.

### Hydrogen production assays

As previously described in Welsh *et al.*^12^, immediately after inoculation of the cultures in 120 mL glass serum vials, sealed with lab-grade butyl rubber stoppers, the headspace of each culture vial was flushed for 5 minutes with 99.99% pure N2. H_2_ concentration was measured using a gas chromatograph containing a pulse discharge helium ionisation detector (model TGA-6791-W-4U-2, Valco Instruments Company Inc) as previously described by Islam et al.^79^. This gas chromatograph was able to detect a wide range of H_2_ concentrations (0.1% – 10% H_2_). Calibration samples of known H_2_ concentration were used to quantify H_2_. The H_2_ concentration within the media-only control vials was measured concurrently to confirm that H_2_ production in isolate samples was biotic.

### Log-ratio of H_2_ producers to H_2_ consumers

The log-ratio of H_2_ producers to H_2_ consumers and flexibles, was calculated as log-ratio of the fraction of H_2_ producers divided by the fraction of H_2_ consumers and flexibles: log2(fraction of H_2_ producers / (fraction of H_2_ consumers + fraction of flexibles) ). R v4.5.1 was used.

### Species richness and species evenness

The species richness and species evenness was calculated for the genomes of all species, only H_2_ producers, only H_2_ consumers, and flexible species. Species richness and species evenness were calculated using the vegan package in R v4.5.1. The Shannon diversity index was computed using the diversity() function with index = “shannon”. Species richness was determined using the specnumber() function, which counts the number of species present in each sample. Pielou’s evenness index was calculated as the ratio of Shannon diversity to the natural logarithm of species richness, following the formula Pielou’s evenness index = Shannon index / ln(species richness). All analyses were performed in R v4.5.1.

### Differential abundance analysis

Differentially abundant species were identified using two tests. First, an unpaired Wilcoxon rank-sum test (Mann-Whitney U test) was computed in R with the function wilcox.test(), a non-parametric method that compares median abundances between the healthy and CD cohort without assuming normality. Second, the package MaAsLin2 (Microbiome Multivariable Association with Linear Models 2) was used to fit multivariable linear models that account for covariates (age and BMI) and compositional effects. One sample without recorded BMI was removed, leaving 85 samples of 47 CD patients and 39 healthy case-controls. To account for the compositional data, the abundance data had been normalised to a total of 1 per sample and a log transform (default) was used in fit_data(). As fixed effects the diagnosis (healthy or CD), age, and BMI were considered (value ∼ diagnosis + age + BMI). No random effects were considered to account for batch effects, as all samples came from the same study.

### Species enrichment analysis

To identify functional groups (H_2_ producers, H_2_ consumers, flexibles, and non-H_2_-cycling species) associated with disease status, we performed a rank-based enrichment analysis using differential abundance results from MaAsLin2. Species were ranked by the MaAsLin2 regression coefficient, i.e. effect size for healthy controls and CD patients, after adjustment for covariates included in the MaAsLin2 model (age and BMI). Species with missing coefficients or duplicated identifiers were excluded prior to ranking. We then applied the GSEA algorithm from the clusterProfiler R package^80^, using the ranked coefficient vector as the input “gene list” and a mapping (“TERM2GENE”) that assigned each species to a metabolic category (i.e.., H_2_ producers, H_2_ consumers, H_2_ flexible taxa, non-cyclers). Enrichment significance was assessed using the default fgsea algorithm and Benjamini–Hochberg multiple-testing correction. Enrichment directionality was interpreted from the normalized enrichment score (NES), with positive NES indicating categories enriched toward the top of the ranked list and negative NES indicating enrichment toward the bottom.

### Biopsy sample collection, culturing and H_2_ measurements

Fresh rectal mucosal biopsy samples were obtained through the endoscopy list at Monash Children’s Hospital (Monash Health; Victoria, Australia) (n = 5) from consenting participants receiving colonoscopy (Ethics approval from the Monash Health Human Research Ethics Committee was granted under HREC/16/MonH/253). One mucosal biopsy sample was taken per participant and used for culturing and subsequent gas production measurements. Participants were classified as Crohn’s disease (CD) or non-inflammatory bowel disease (non-IBD) controls based on standard clinical, endoscopic and histopathological criteria. Biopsies were immediately placed into an anaerobic environment and transported using anaerobic jars with an Oxoid™ AnaeroGen™ 2.5L Sachet. All processing steps were performed under anaerobic conditions. Samples were homogenised using 130 µL sterile, pre-reduced PBS for 5 minutes. Initial bacterial load was quantified by plating 10 µL aliquots of the homogenate in technical triplicates on pre-reduced YCFA agar plates for colony-forming unit (CFU) enumeration, following serial dilution in sterile, pre-reduced PBS. 200 µL of homogenate was inoculated into 30 mL pre-reduced YCFA broth, in triplicate independent cultures in 120 mL serum vials. Cultures were flushed with pure N_2_ for 6 minutes and incubated anaerobically at 37 °C. 2 mL headspace gas samples were collected at 24 h and stored in 3 mL glass exetainers. H_2_ concentrations were measured by gas chromatography using a thermal conductivity detector, and production was calculated as the change over time.

### Abundance-weighted metabolic versatility of fermentation routes

As before, metagenomic stool samples from CD patients and healthy case-controls from He *et al.*^70^ were used. The relative abundances of H_2_ producing species for healthy controls and CD patients were normalized within each sample to a sum of 1, disregarding all non-H_2_-producers. For each H_2_ producer species, the number of predicted fermentation end products was obtained from the previous analysis of H_2_ producer capabilities. Sample-level fermentation versatility was then calculated as an abundance-weighted sum across species (i.e., the normalized abundance of each species multiplied by its number of predicted fermentation products, summed per sample). Group means were computed for the healthy and CD cohort, and 95% confidence intervals were estimated using the *t*-distribution, shown as bar plot (mean ± 95% CI) with jittered points representing individual samples. Differences between conditions were evaluated using the Wilcoxon rank-sum test.

### Shifts in fermentation guild composition

As before, metagenomic stool samples from CD patients and healthy case-controls from He *et al.*^70^ were used and the relative abundances of H_2_ producing species were normalized within each sample to a sum of 1, disregarding all non-H_2_-producers. To assess systematic changes in fermentation routes, H_2_ producer species were then grouped into functional “fermentation guilds” defined by identical sets of predicted fermentation products (i.e., identical binary fermentation profiles). Each species was assigned to a guild and then guild-level relative abundances were computed per sample by summing up the normalized abundances of all species belonging to the same guild. Mean guild abundances were calculated for healthy controls and CD patients. The guilds were then filtered to retain those with mean relative abundance > 5% (with all H_2_ producers summing up to 100%) in at least one condition and an absolute mean difference > 5% between the healthy and the CD cohort. Mean abundances were plotted as horizontal bars for healthy controls and CD patients, with jittered points showing per-sample guild abundances. For each retained guild, differences in abundance between healthy and CD were tested using Wilcoxon rank-sum tests (with exact = FALSE to accommodate tied values arising from zeros).

### H_2_S detection via methylene blue assay

H_2_S in bacterial culture supernatants was quantified using a methylene blue colorimetric assay. *Sulfide colour reagent* was prepared by dissolving 0.4 g *N,N’*-dimethyl-1,4-phenylenediamine sulfate in ∼25 mL Milli-Q water, adding 50 mL concentrated HCl, and filling to 100 mL with Milli-Q water. *Sulfide catalyst solution* was prepared by dissolving 1.6 g FeCl_3_ 6H_2_O in ∼40 mL Milli-Q water, adding 50 mL concentrated HCl, and filling to 100 mL with Milli-Q water. Immediately before use, equal volumes of colour reagent and catalyst solution were mixed to generate the *sulfide reagent*. *Zinc acetate preservative* was prepared by dissolving 0.261 g zinc acetate in 25 mL Milli-Q water.

Cultures of bacterial isolates were grown anaerobically in 10 mL YCFA broth in 120 mL bottles for 12 h under either high H_2_ (100% H_2_ headspace) or low H_2_ (100% N_2_ headspace) conditions. Triplicates of the following conditions were analysed: cell-free medium as a negative control, *asr*-negative *Bacteroides uniformis*, *asr*-positive *Enterocloster clostridioformis*, and *asr*-positive *Clostridium symbiosum*. 200 μM sodium sulfide (Na_2_S) standard was included as a positive control. To preserve dissolved sulfide, zinc acetate was anaerobically added to cultures at 0.1 mL per 2 mL sample. Sulfide reagent was then added at 0.1 mL per 2 mL sample or standard and mixed. Samples were transferred to 1.5 mL centrifuge tubes and cells were pelleted by centrifugation for 2 min at 14,800 rpm. 1 mL clarified supernatants were transferred to cuvettes and absorbance was measured at 670 nm as a qualitative measure of H_2_S concentrations.

## Acknowledgments

We thank Matthias Hülsmann for insightful discussions on redox balancing, Oliver Schmidt for helpful discussions of fermentation, as well as Tom Watts, Bob Leung, Zahra Islam, and current and previous members of the Greening group for helpful discussions and questions. We also extend our gratitude to Jovita Quach who transported the biopsy samples from the endoscopy theatre to the laboratory.

We acknowledge the Australian Microbiome Culture Collection, from which we obtained the isolates CC01283 (*A. rectalis*), CC01011 (*B. uniforms*), CC00689 (*B. fusiformis*), CC00294 (*B. plebeius*), CC00281 (*C. innocuum*), CC01278 (*E. clostridioformis*), CC00403 (*H. effluvii*), and CC00921 (*C. symbiosum*).

## Author contributions

Conceptualization: AS, CG

Methodology: AS, CG

Investigation: AS, CW, LL, YK, EG

Visualization: AS

Funding acquisition: CG, AS

Project administration: AS Supervision: CG

Writing – original draft: AS

Writing – review & editing: AS, CG, and all other co-authors

